# Transcription-induced coacervation accelerates and sensitizes cell-free biosensing

**DOI:** 10.64898/2026.07.02.736143

**Authors:** Siyuan Feng, Rebecca Rasmussen, Antonio Garcia, Lauren Clark, Samanvaya Srivastava, Julius B. Lucks

**Author notes:** To whom correspondence should be addressed. Emails: Samanvaya Srivastava,; Julius B. Lucks,.

## Abstract

Cell-free biosensors leverage *in vitro* gene expression reactions to detect chemicals. While inexpensive, modular, and distributable, these platforms are constrained by slow readouts at ambient temperatures, precluding practical field operation. In cells, phase separation accelerates biochemical reactions; however, recapitulating these gains *in vitro* has remained challenging for complex biochemistries. Here, we report the first self-assembling coacervate system that accelerates *in vitro* transcription. Prepared by simple mixing, coacervation with spermine and polyacrylic acid occurs dynamically in response to NTP consumption and co-localizes DNA templates and RNA polymerase to accelerate transcription, mimicking intracellular phenomena. We exploit this discovery to accelerate the cell-free biosensing of six ligands, demonstrating that coacervation can preserve platform modularity, improve sensitivity, retain lyophilization compatibility, function in field matrices, and reduce ambient-temperature time-to-signal by hours. This work contributes to a growing understanding of phase separation in biology and advances the use of membrane-less organization for real-world applications.

## INTRODUCTION

Over the past two decades, *in vitro* cell-free biosensors have been developed to sense a broad range of analytes, including antibiotics [1], metal ions [1, 2], viruses [3, 4], miRNA [5], hormones [6], neurotransmitters [7, 8], personal health products [1], herbicides [9], stimulants [10], toxins [11–13], and disease markers [1, 10, 14–16]. These systems repurpose molecular machinery found in cells and typically consist of a sensing element, such as a ligand-responsive DNA, RNA, or protein, and a reporter element such as a fluorescent protein, with the elements coupled through molecular chemistries including transcription, translation, or cleavage [17, 18]. Cell-free biosensors are modular and can be reconfigured to sense many compounds [1, 14, 15], can be interfaced with molecular circuits that improve performance with logic, feedback, and signal amplification [4, 5, 19], optimized with computational design [2, 5, 20], lyophilized for use in the field [1, 11, 14, 16, 19] and made usable to non-expert individuals in low-resource settings [13], and have thus found broad relevance in environmental and human health applications [17, 18, 21].

While promising, a major challenge constraining the real-world utility of cell-free biosensors is their long time-to-signal at ambient temperatures, requiring operation inside 37 °C incubators that are costly and inconvenient [22]. These incubators are also bulky and require battery charging, adding both an infrastructure requirement and a failure mode in the field. Even at optimal incubation temperatures, the fastest field-deployable cell-free biosensors can take hours to turn on [17], due to slow sensing mechanisms, lyophilization effects, and complex sample matrices. Other approaches to improve sensing kinetics have included strand displacement circuits [19] and CRISPR-Cas-based schemes [23]; these strategies often involve complex set-ups, expensive modified oligonucleotides, additional purified proteins, and still require incubation at 37 °C for optimal performance [19, 23, 24].

Here, we develop a simple, low-cost approach to accelerate ambient-temperature cell-free biosensing by leveraging liquid-liquid phase separation, which cells routinely use to accelerate slow biochemical reactions [25]. Liquid-liquid phase separation, also known as condensate formation, is used to co-localize reactants [25], sequester inhibitors [25], and generate favorable environments with appropriate redox conditions, pH, and water [26–28], facilitating both intracellular protein and nucleic acid chemistries. Coacervates, aqueous liquid-like droplets formed by charged multivalent polymers, have emerged as a model for these phase-separated systems [29, 30], with engineered coacervates in cells demonstrating spatial control over transcription [31], enhanced gene expression [32, 33], and accelerated enzymatic conversion [34]. Coacervation applied to *in vitro* systems has also shown promising results, especially in accelerating simple biochemical reactions, including actin assembly and ribozyme function [35, 36], concentrating nucleoside triphosphates (NTPs) and magnesium [37], and supporting the folding and function of key biosensor parts such as ribozymes and aptamers [38–40]. Importantly, coacervation can be induced simply by mixing commercial, low-cost polymers readily available at scale. We thus hypothesized that coacervates could be harnessed to accelerate reaction kinetics in cell-free systems and enhance biosensing performance, especially at ambient temperatures.

While prior research has identified *in vitro* coacervates that enhance DNA hybridization [41] and hemin peroxidative activity [42] to improve molecular biosensing, these approaches are limited to analytes that can be directly recognized by DNA and were not assessed in field formats. Other efforts to couple biosensing with phase-separated systems have generated stimulus-responsive coacervates but have not improved sensing performance [43–45]. In fact, efforts to couple coacervates with cell-free systems have consistently yielded lower reaction efficiencies [46–49] or required assembly via microfluidics, which is not easily deployed in the field [50]. Therefore, we sought to develop a spontaneously assembling coacervate system to enhance *in vitro* transcription (IVT), a core platform upon which most cell-free technologies are built [17, 18]. Successful coupling of coacervates to a cell-free system would maintain key strengths, including modularity and compatibility with lyophilization, while achieving accelerated sensing of a broad range of analytes at ambient temperatures.

Here, we achieve this goal by engineering a coacervate system that accelerates the kinetics of our previously developed RNA Output Sensors Activated by Ligand INDuction (ROSALIND) cell-free biosensing platform, which uses transcription factor-regulated IVT of fluorescent RNA aptamers to detect target compounds [1]. We first investigate three coacervate compositions and show that spermine-polyacrylic acid (PAA) coacervates dramatically accelerate IVT such that the kinetics of room-temperature IVT with coacervates exceed those of control IVT reactions at 37 °C. We discover that coacervation with spermine and PAA is dynamic and is induced by IVT through NTP hydrolysis. We further show that this system accelerates transcription through the co-concentration of DNA and T7 RNA polymerase (T7 RNAP) at the surface of the coacervate droplets. We then establish enhanced, modular biosensing at room temperature for six unique ligands, including antibiotic analogs, metals, and plant metabolites, showing that coacervates can reduce time to signal by hours, often with improved detection sensitivity. Lastly, we demonstrate that coacervates are amenable to single-pot lyophilization with biosensor parts and that coacervate-mediated enhancement of biosensing is preserved upon rehydration of lyophilized reactions with unprocessed lake water matrices. Taken together, our results show that self-assembling transcription-induced coacervates can be used to accelerate and sensitize cell-free biosensing to meet real-world needs.

## RESULTS

### Spermine-PAA coacervates increase the signal kinetics of T7 RNAP *in vitro* transcription

We first sought to identify complex coacervate compositions that self-assemble by mixing and improve IVT by T7 RNAP. Although other coacervate systems have been developed to interface with cell-free transcriptional or translational processes [46, 48, 50–52], to our knowledge, no self-assembling, phase-separated system has been reported to enhance these *in vitro* processes. To assess how polyelectrolyte identity affects compatibility with IVT, we investigated coacervate systems assembled by mixing the polyanion PAA with three polycations: polyallyldiammonium chloride (PDADMAC), spermine, and spermidine. We chose PAA for its broad use and biocompatibility, as demonstrated in other applications [39, 53, 54], while choosing PDADMAC, spermine, and spermidine because of their previous use in generating coacervates that support non-enzymatic RNA synthesis [55], catalyze ribozyme assembly [39], and concentrate solutes [56], respectively.

To identify polyelectrolyte pairs that undergo coacervation in IVT buffer, we mixed PAA with each polycation in buffer (**Supplementary Table 1**), and quantified the mixture’s turbidity to assess phase separation [53, 57]. Mixing PAA with either PDADMAC or spermine in IVT buffer generated markedly turbid solutions, suggesting that these polyelectrolyte pairs phase-separate robustly in the buffer (**Fig. 1A**). We also confirmed that both PDADMAC-PAA and spermine-PAA systems form coacervates in IVT buffer using DIC microscopy (**Fig. 1B-C**). To investigate compatibility with IVT reactions, we introduced PAA with either PDADMAC or spermine into IVT reactions containing a reporter template encoding the fluorescence-activating RNA aptamer, three-way junction dimeric Broccoli (3WJdB) [58] (**Fig. 1D**). Transcription reactions were supplemented with the 3WJdB-binding dye DFHBI-1T and monitored for the production of fluorescence, indicative of 3WJdB RNA synthesis and folding, at 37 °C. Excitingly, we discovered that, while PDADMAC-PAA coacervates inhibit IVT (**Fig. 1E**), spermine-PAA coacervates accelerate IVT signal kinetics (**Fig. 1F**).

**Figure 1:**
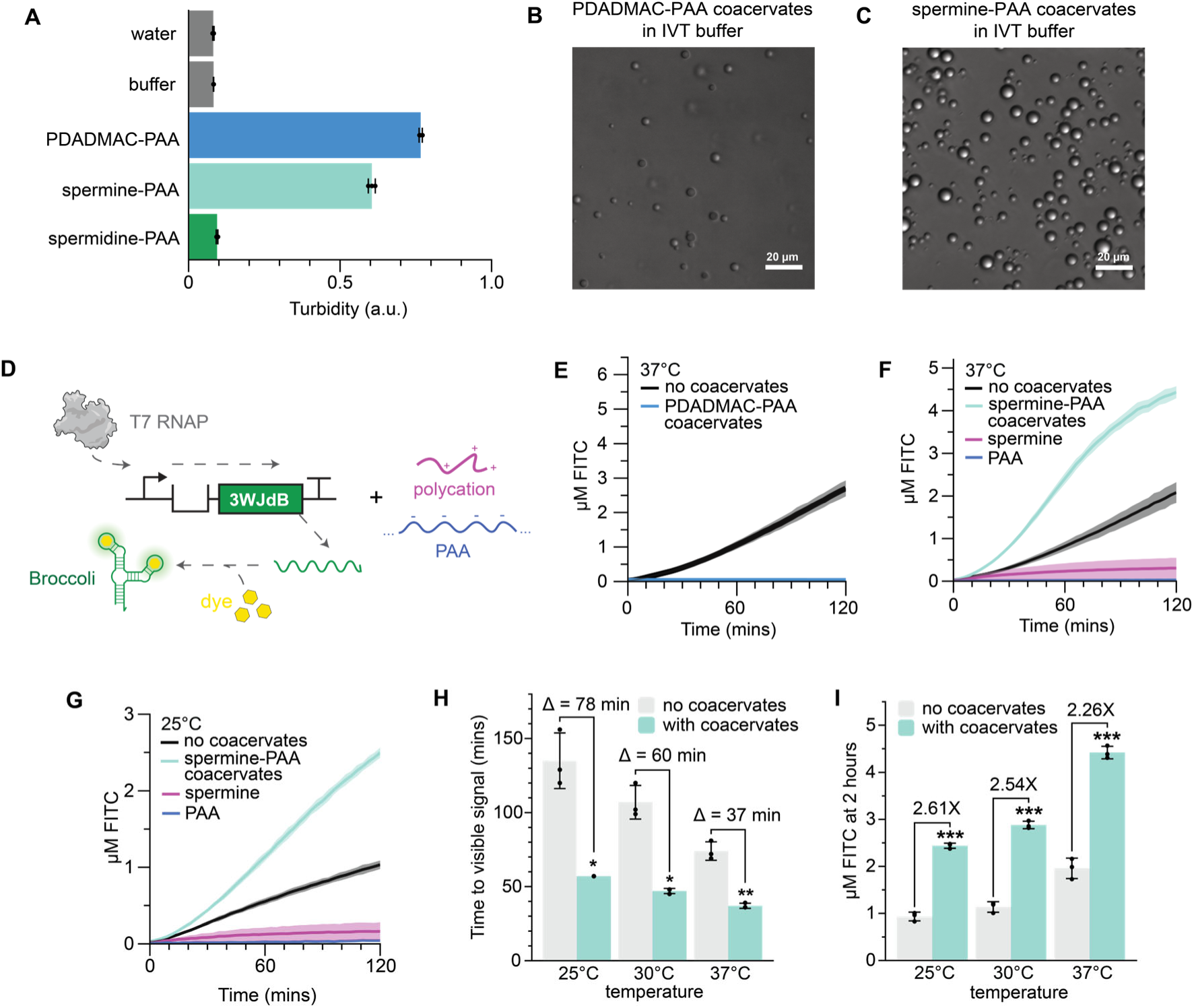
Self-assembling spermine-PAA coacervates enhance IVT signal kinetics. **A-C**, Mixing PDADMAC or spermine in IVT buffer with PAA generates phase separation *in situ*, as indicated by the increase in turbidity (**A**) and visualized by differential interference contrast (DIC) microscopy (**B,C**) immediately after mixing. Mixing spermidine with PAA resulted in minimal change to solution turbidity, suggesting weak phase separation in IVT buffer. **D,** Schematic of T7 RNAP-driven *in vitro* transcription of 3WJdB, with added polyelectrolytes. **E**, Addition of PDADMAC-PAA coacervates to IVT reactions inhibits IVT signal at 37 °C. **F-G,** Addition of spermine-PAA coacervates improves IVT signal kinetics at 37 °C (**F**) and 25 °C (**G**), while the addition of spermine or PAA alone does not. **H-I,** Addition of spermine-PAA coacervates reduces time to visible signal (**H**) and increases fluorescence after 2 hours (**I**) for reactions run at 25, 30, and 37 °C. A two-tailed, heteroscedastic Student’s *t*-test was used to compare samples with and without coacervates at each temperature. The data shown in **A,H,I** are *n* = 3 independent replicates, each plotted as a point, with the bar heights representing the average over these replicates and error bars indicating the standard deviation. For the data in **E**,**F,G**, the shading indicates the average value of n=3 independent biological replicates +/- standard deviation. The *P* value range in (**H,I**) is indicated by asterisks (\*\*\**P* < 0.001, \*\**P* = 0.001–0.01, \**P* = 0.01–0.05, ns otherwise). Time to visible signal is defined as the time to 1 µM FITC, based on 3WJdB fluorescence visualized in previous work [1]. The polyelectrolyte concentrations and the DNA sequences used in each experiment are listed in **Supplementary Table 1** and **Supplementary Data 1**. NTPs and TIPP are included in the IVT reaction but are not shown in the schematic in **D**. Raw fluorescence values were standardized to µM FITC.

To demonstrate that the enhancement in transcriptional signal is due to the presence of coacervates, we tested the effect of adding spermine or PAA alone, and found that individually, these polymers suppress transcription, indicating that their complexation is required to enable transcription (**Fig. 1F**). To investigate whether spermine-PAA coacervates can enhance readout kinetics at other temperatures, we assembled IVT reactions with spermine and PAA, incubated at 25, 30 and 37 °C, and monitored 3WJdB fluorescence. At each temperature, we found that adding spermine-PAA coacervates led to faster transcription kinetics (**Fig. 1G-H**) and higher signal after two hours (**Fig. 1I**), with greater fold-changes at cooler temperatures. Notably, coacervate-enhanced IVT kinetics at 25 °C were faster than reactions run at 37 °C without coacervates (**Fig. 1H**).

Together, these results show that the chemical identity of polyelectrolytes can modulate the compatibility of coacervates with IVT, and that self-assembling spermine-PAA coacervates can enhance IVT signal kinetics, particularly at ambient temperatures.

### Spermine-PAA coacervate formation is dynamically induced by *in vitro* transcription

Although we found that spermine and PAA form droplets immediately upon mixing in transcription buffer (**Fig. 1C**), we also observed that, when assembled in full IVT reactions, coacervate formation was not immediate (**Fig. 2A-B**). We reasoned that one of the IVT reaction components could be inhibiting immediate coacervation. To identify this component, we sequentially omitted reaction components from otherwise complete IVT reactions and discovered that NTPs inhibit coacervate formation (**Fig. 2C**). Partial removal of NTPs was sufficient to result in immediately turbid solutions after mixing (**Fig. 2D**), with observable droplets under microscopy (**Fig. 2E**). To assess whether the RNA generated from NTPs also contributes to coacervation, we added purified RNA to IVT reactions containing spermine and PAA, without T7 RNAP to preclude NTP hydrolysis, and found that the addition of RNA had no measurable impact on turbidity (**Supplementary Fig. 2**).

**Figure 2:**
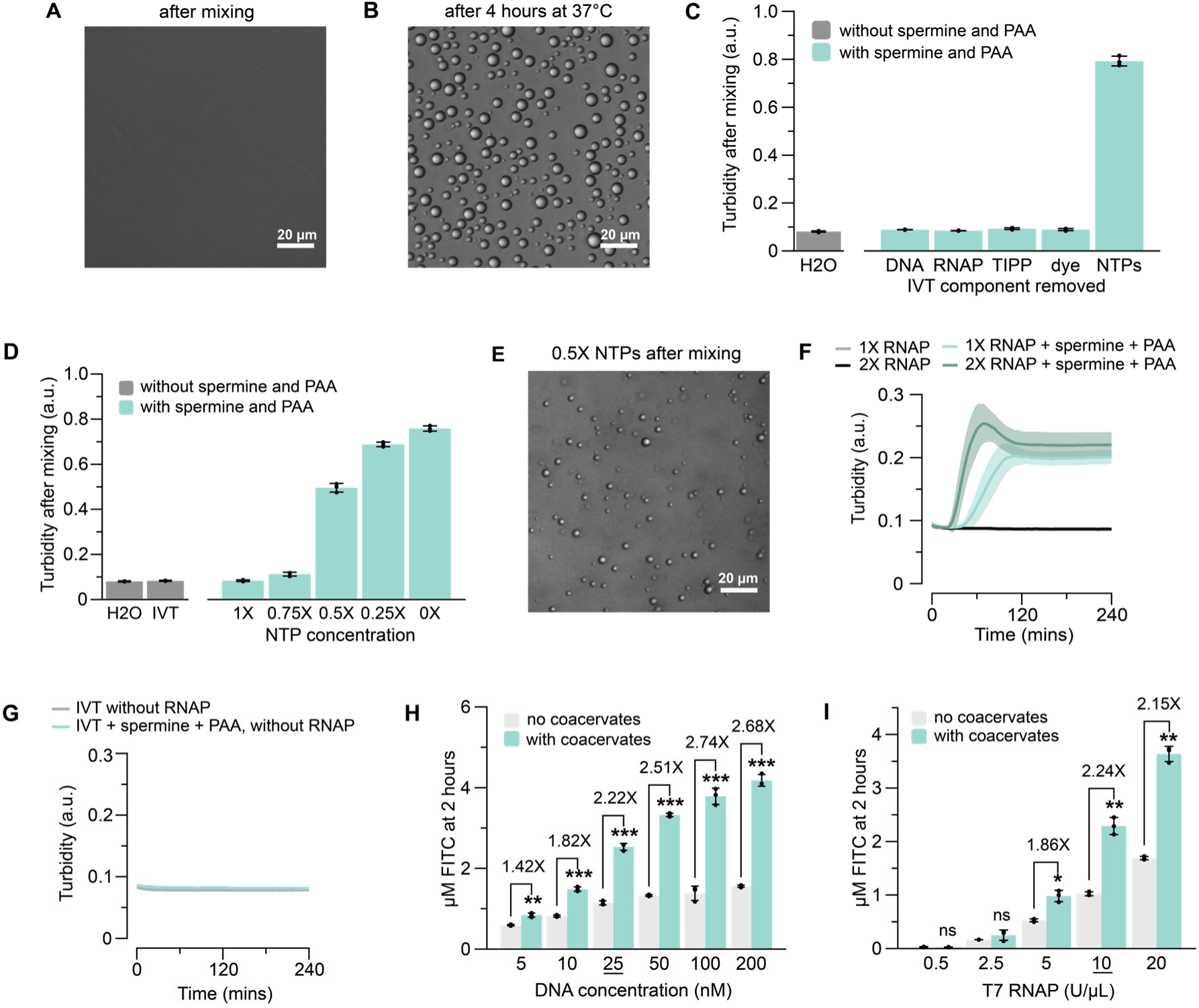
Spermine-PAA coacervate formation is dynamically induced by IVT through NTP removal. **A-B**, Coacervate formation is not immediate upon spermine and PAA addition to IVT reactions. DIC images show that coacervates are not present immediately after mixing (**A**) but are observable after 4-hour incubation at 37 °C (**B**). **C**, Reaction components were sequentially omitted from IVT reactions to elucidate the components that inhibit coacervate formation. IVT reactions with spermine and PAA, assembled without NTPs, become turbid immediately after mixing. **D**, Turbidity measurements of IVT reactions without T7 RNAP show that reducing the NTP concentration is sufficient to induce turbid solutions immediately after mixing. T7 RNAP was removed to preclude additional NTP consumption. The turbidity measurements in **C**,**D** were taken on a plate reader set to room temperature after reaction assembly on ice. **E,** Representative image taken of an IVT reaction with 0.5X less NTPs without T7 RNAP immediately after mixing. **F**, Turbidity measurements over time of IVT reactions using 1X or 2X RNAP at 37 °C, with or without spermine and PAA, show that increasing turbidity, indicative of coacervation, occurs with *in vitro* transcription with spermine and PAA. Using more RNAP expedites the onset of measurable turbidity. **G**, IVT reactions assembled with spermine and PAA, but without T7 RNAP, do not become turbid while incubated at 37°C, indicating that time and incubation alone are insufficient to induce phase separation. **H-I**, Coacervates enhance IVT signal across a range of DNA and RNAP concentrations. IVT reactions assembled with spermine-PAA coacervates demonstrate higher signal after 2 hours of incubation at room temperature for several DNA (**H**) and RNAP (**I**) concentrations. Enhancement increases with increasing DNA or RNAP concentrations. A two-tailed, heteroscedastic Student’s *t*-test was used to compare samples with and without coacervates at each condition. The underlined values correspond to the 1X DNA and 1X T7 RNAP concentrations in a typical IVT reaction. The data shown in **C,D,H,I** are *n* = 3 independent biological replicates, each plotted as a point, with the bar heights representing the average over these replicates and error bars indicating the standard deviation. For the data in **F,G**, the shading indicates the average value of n=3 independent biological replicates +/- standard deviation. The *P* value range in **H,I** is indicated by asterisks (\*\*\**P* < 0.001, \*\**P* = 0.001–0.01, \**P* = 0.01–0.05, ns otherwise). The DNA sequences used in each experiment are listed in **Supplementary Data 1**. Raw fluorescence values were standardized to µM FITC.

Based on these findings, we hypothesized that coacervation is triggered by ongoing transcription, primarily through NTP consumption. This hypothesis is consistent with prior work showing that biochemical reactions [59, 60], including NTP consumption [45, 61], influence the formation and stabilization of coacervate systems, as well as the known role of ATP in solubilizing phase-separated compartments [62]. To verify this hypothesis, we assembled IVT reactions with spermine and PAA and quantified turbidity over time (**Fig. 2F**). We discovered that the onset of measurable turbidity occurs around 30 mins after assembly. In addition, we found that accelerating transcription by including more RNAP enzyme expedites the onset of measurable turbidity (**Fig. 2F**). To determine whether coacervate formation is linked to transcription, we assembled IVT reactions with spermine and PAA, but without T7 RNAP, and observed that they do not become turbid, suggesting that transcription is required to induce phase separation (**Fig. 2G**). To confirm that *in vitro* transcription modulates coacervate formation, we imaged IVT reactions at increasing RNAP concentrations and observed that enhancing IVT with increased RNAP yields more and larger droplets, which may reflect coalescence as the number of coacervate droplets increases (**Supplementary Fig. 3**).

To understand how this mechanism affects the coacervate-mediated enhancement of signal readout, we assembled IVT reactions with increasing concentrations of DNA or RNAP. We found that coacervates enhance IVT signal across a range of DNA and RNAP concentrations, with greater enhancement at higher DNA or RNAP concentrations (**Fig. 2H-I**), which we hypothesize is due to an earlier onset of coacervation under stronger IVT conditions.

Together, these results indicate that NTP consumption governs the formation of spermine-PAA coacervates in IVT reaction mixtures, with enhanced IVT leading to earlier coacervation and greater coacervate-mediated enhancement.

### Spermine-PAA coacervates accelerate transcription by co-concentrating DNA and RNAP

Having uncovered how spermine-PAA coacervates self-assemble in IVT reactions, we next sought to investigate the mechanisms by which these coacervates enhance IVT signal readout. 3WJdB signal generation kinetics depend on both transcription rate, which determines how much 3WJdB RNA is synthesized, and non-transcriptional processes, including DFHBI-1T dye binding and fluorescence [63, 64]. Based on previous work showing that coacervates can enhance catalytic processes [30, 65], we hypothesized that spermine-PAA coacervates increase IVT signal by accelerating RNAP transcription, rather than post-transcriptional processes. To validate this hypothesis, we quantified the amount of RNA generated in room temperature IVT reactions using a Qubit quantification assay (**Fig. 3A**) and found that coacervates result in a 2-fold increase in the amount of RNA generated after 2 hours of incubation, matching the roughly 2-fold improvement observed in our fluorescent readout studies (**Fig. 1I**). We verified this finding using urea-PAGE gel electrophoresis as a complementary approach (**Supplementary Fig. 4**).

**Figure 3:**
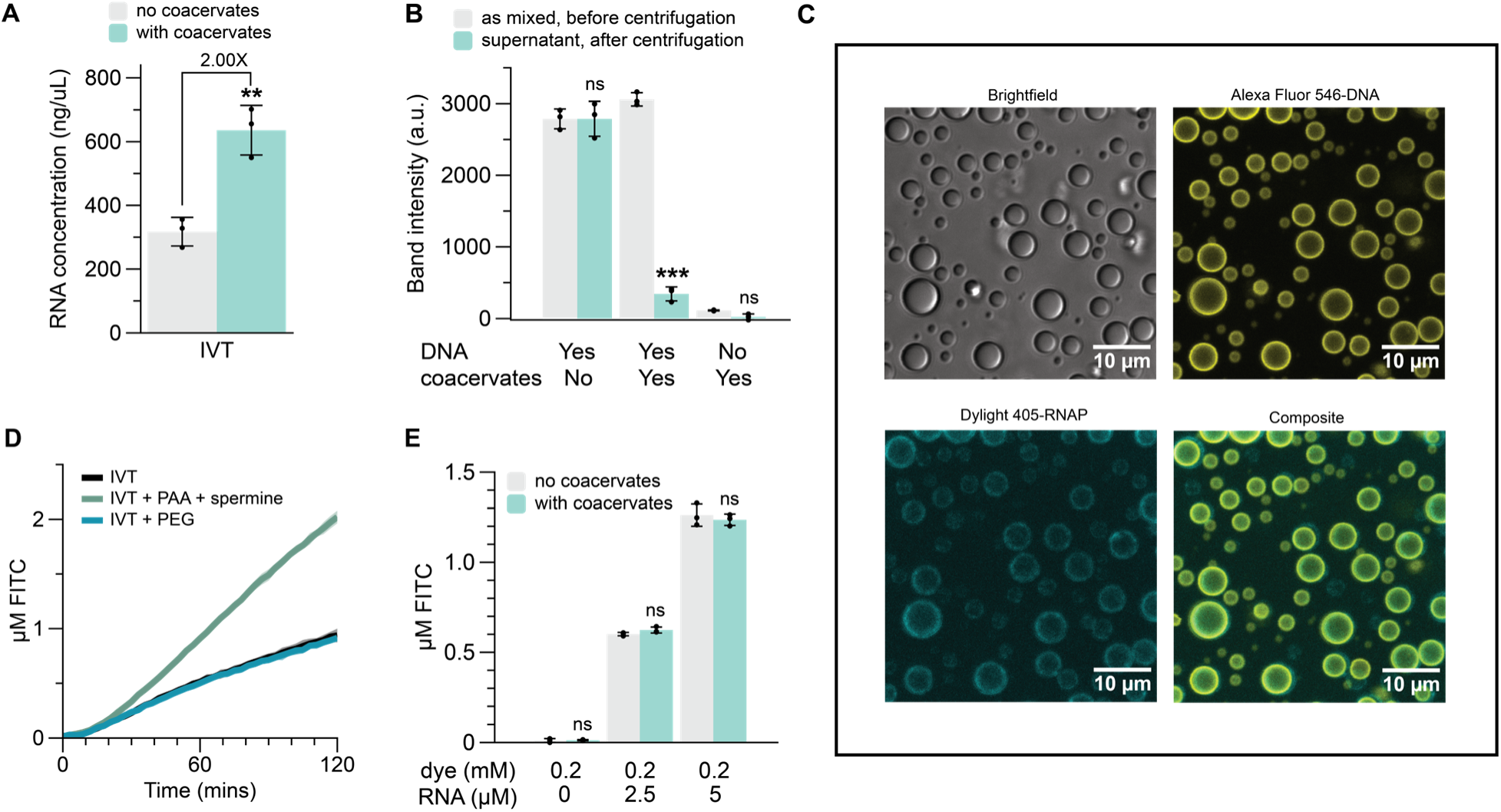
Spermine-PAA coacervates accelerate IVT by co-concentrating DNA and RNAP. **A**, IVT quantification by RNA Qubit assay after RNA isolation shows that IVT reactions with coacervates generate 2-fold more RNA than reactions without coacervates. IVT reactions were incubated for two hours at room temperature (∼25 °C). A two-tailed, heteroscedastic Student’s *t*-test was used to compare samples with and without coacervates. **B,** Quantification of DNA in transcription buffer with coacervates before and after centrifugation to pellet the coacervate phase. A drop in DNA concentration in the post-centrifugation supernatant indicates that DNA is sequestered by the coacervate phase. The DNA was quantified using gel electrophoresis (**Supplementary Fig. 6**), with the band intensities measured using ImageJ/FIJI. A two-tailed, paired Student’s *t*-test was used to compare samples before and after centrifugation. **C,** Pseudo-brightfield and confocal microscopy images with Alexa Fluor 546-labeled DNA (yellow) and DyLight 405-labeled T7 RNAP (cyan), showing that these molecules co-localize strongly to the coacervate interface. Samples for microscopy were prepared by mixing the fluorescent molecules with transcription buffer, spermine, and PAA. **D,** Kinetic traces of IVT reactions containing an equiweight concentration of molecular crowder PEG to replace spermine and PAA, indicating that spermine and PAA increase transcription through mechanisms beyond volume crowding generated simply by their addition to the reaction mixture. Reactions were run at room temperature (∼25 °C). **E,** Fluorescence measurements of mixtures containing coacervates with different concentrations of purified 3WJdB RNA and its cognate dye DFHBI-1T, demonstrating that coacervates do not non-specifically activate DFHBI-1T in the absence of 3WJdB RNA or enhance 3WJdB RNA fluorescence. Measurements were taken after two hours of incubation at room temperature (∼25 °C). A two-tailed, heteroscedastic Student’s *t*-test was used to compare samples with and without coacervates. The data shown in **A,B,E** are *n* = 3 independent biological replicates, each plotted as a point, with the bar heights representing the average over these replicates and error bars indicating the standard deviation. For the data in **D**, the shading indicates the average value of n=3 independent biological replicates +/- standard deviation. The *P* value range in **A,B,E** is indicated by asterisks (\*\*\**P* < 0.001, \*\**P* = 0.001–0.01, \**P* = 0.01–0.05, ns otherwise). The DNA and RNA sequences used in each experiment are listed in **Supplementary Data 1**. Raw fluorescence values were standardized to µM FITC.

To explain how coacervates increase RNA production, we hypothesized that they enrich key IVT components, thereby increasing effective reaction concentrations, consistent with earlier reports for other biological processes [36, 54, 61, 66]. To test this hypothesis, we mixed 25 nM DNA with spermine-PAA coacervates in IVT buffer and separated the dilute supernatant from the complex coacervate phase by centrifugation, using turbidity measurements to verify the complete removal of coacervate droplets (**Supplementary Fig. 5**). We then measured the concentration of DNA in the dilute supernatant phase using gel electrophoresis, which revealed that more than 85% of the DNA was sequestered in the dense coacervate phase (**Fig. 3B** and **Supplementary Fig. 6**). We estimate that the dense phase constitutes less than 5% of the total volume (**Supplementary Fig. 7**), which corresponds to a DNA concentration that is, on average, 17-fold higher in the coacervate phase. Additionally, confocal microscopy using an IVT template DNA labeled with Alexa Fluor 546 revealed that the fluorophore-labeled DNA localizes strongly to the coacervate phase (**Supplementary Fig. 8**), most notably at the coacervate interface, a pattern of localization that has been observed for other biomolecules [36, 65, 67–69]. Similarly, confocal microscopy experiments with T7 RNAP labeled with DyLight 405 fluorophore revealed that fluorophore-labeled T7 RNAP also concentrates strongly to this location (**Supplementary Fig. 9**) and co-localizes with the DNA (**Fig. 3C**). These results suggest that spermine-PAA coacervates enhance IVT signal by co-concentrating DNA and RNAP, thereby accelerating transcription kinetics.

Macromolecular crowding agents such as poly(ethylene glycol) (PEG) have also been used to occupy reaction volume and enhance nucleic acid synthesis [70, 71]. To test whether simple crowding by spermine and PAA, independent of coacervation, could account for the observed IVT enhancement in our system, we added an equiweight concentration of PEG to replace both spermine and PAA and found that transcription was not enhanced at this PEG concentration (**Fig. 3D**). We also assembled reactions that replace only PAA with PEG and discovered that introducing spermine with PEG inhibits transcription (**Supplementary Fig. 10**). These results suggest that spermine and PAA enhance IVT signal through mechanisms beyond macromolecular crowding generated simply by their addition to the reaction mixture.

Some fluorophores are also reported to exhibit enhanced quantum yield in coacervates [41]. To investigate whether spermine-PAA coacervates enhance signal readout by increasing 3WJdB-DFHBI-1T fluorescence, we added purified 3WJdB RNA to DFHBI-1T dye in transcription buffer, in mixtures with or without spermine and PAA, and found that adding spermine and PAA to purified 3WJdB resulted in unchanged fluorescence (**Fig. 3E**). We also added spermine and PAA to DFHBI-1T alone and noted that coacervates do not increase signal readout from non-specific activation of DFHBI-1T (**Fig. 3E**). These results suggest that spermine-PAA coacervates do not enhance IVT signal by increasing the fluorescent quantum yield of 3WJdB-DFHBI-1T, and are consistent with prior work demonstrating the same for the stabilized dimeric Broccoli aptamer (sdB) in PDADMAC-RNA coacervates [55].

Together, these results show that spermine-PAA coacervates accelerate IVT reaction kinetics by co-concentrating DNA and RNAP.

### Coacervates accelerate transcriptional biosensing and preserve platform modularity

After demonstrating that spermine-PAA coacervates accelerate IVT, we sought to explore whether this strategy could be extended to enhance the detection of chemical targets in transcriptional biosensing reactions that build on IVT. Our previous work established the ROSALIND system, which incorporates allosteric transcription factors (aTFs) to regulate the IVT synthesis of fluorescent RNA reporters to detect chemical targets [1] (**Fig. 4A**). In ROSALIND, repressor aTF proteins bind operator sequences located downstream of T7 RNAP promoters within DNA reporter templates encoding for 3WJdB, thereby repressing aptamer transcription by T7 RNAP. In the presence of their cognate ligand, these aTFs unbind the operator sequence and permit reporter transcription, generating a fluorescent readout.

**Figure 4:**
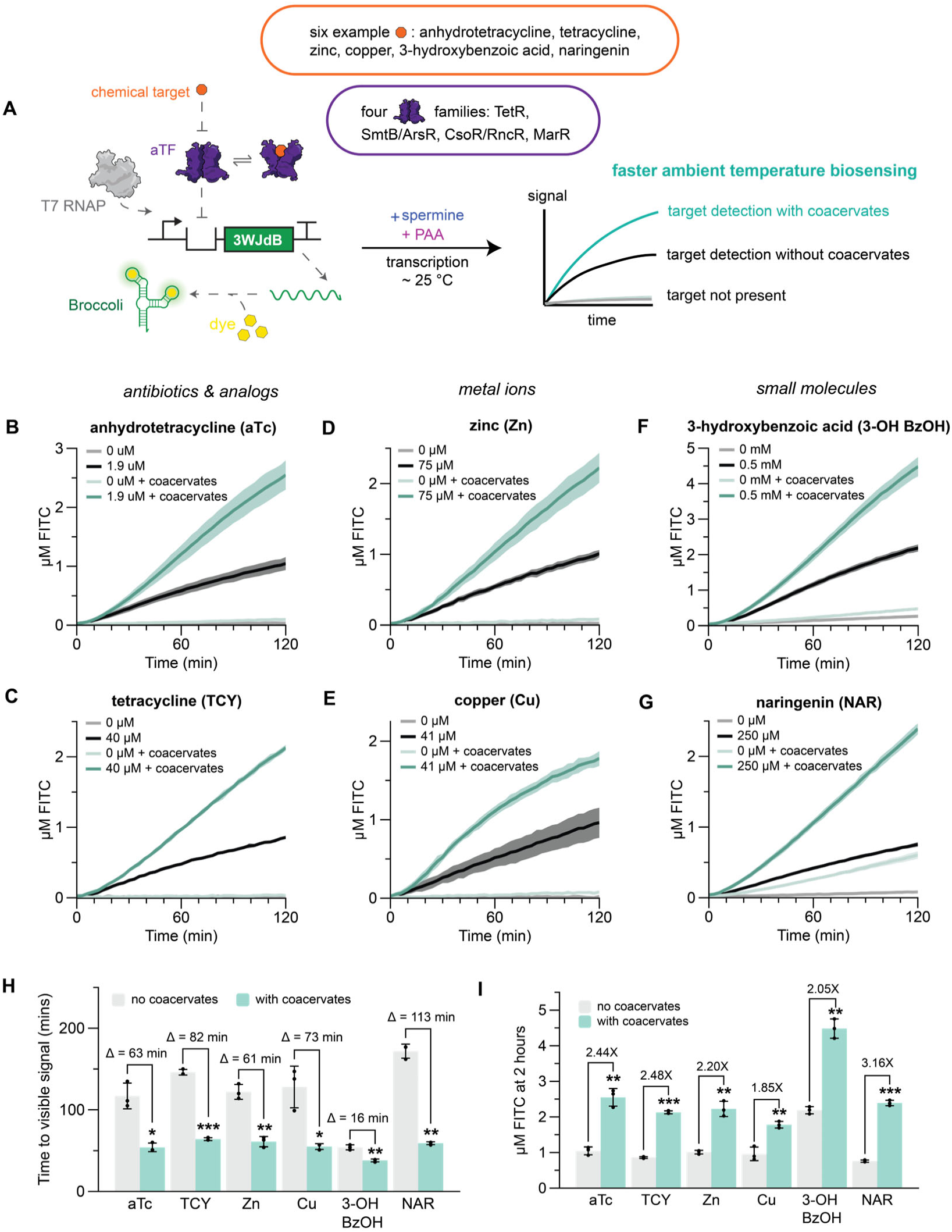
Coacervates accelerate transcriptional biosensing of multiple chemical targets at ambient temperature. **A,** Spermine-PAA coacervates support the complex biochemistry required for transcriptional biosensing, including aTF repression and induction, RNAP transcription, and RNA aptamer folding. Coacervates can leverage multiple aTF families, accelerating the sensing of diverse ligands, including (**B,C**) anhydrotetracycline and tetracycline sensing with TetR, (**D**) zinc sensing with SmtB, (**E**) copper sensing with CsoR, (**F**) 3-hydroxybenzoic acid sensing with MobR, and (**G**) naringenin sensing with TtgR. **H,** Coacervates reduce time to visible signal by an average of 68 minutes and **I,** increase fluorescence at 2 hours by approximately 2-fold for each ligand. Each reaction contains the indicated ligand concentration (as prepared in **Supplementary Table 3**) or a laboratory-grade water control. DNA template and aTF concentrations used in each reaction are listed in **Supplementary Table 3**, and the DNA and protein sequences are listed in **Supplementary Data 1**. All reactions were run at room temperature (∼25 °C). For the data in **B** – **G**, the shading indicates the average value of n=3 independent biological replicates +/- standard deviation. The data shown in **H** and **I** summarize the kinetic traces shown in **B** – **G,** with bar heights representing the average of n=3 independent replicates, each plotted as a point, and error bars indicating the standard deviation. aTc, TCY, 3-OH BzOH, and NAR are abbreviations for anhydrotetracycline, tetracycline, 3-hydroxybenzoic acid, and naringenin. A two-tailed, heteroscedastic Student’s *t*-test was used to compare the time to signal between conditions with and without coacervates. The *P* value range in **H,I** is indicated by asterisks (\*\*\**P* < 0.001, \*\**P* = 0.001–0.01, \**P* = 0.01–0.05, ns otherwise). Time to visible signal is defined as the time to 1 µM FITC, based on 3WJdB fluorescence visualized in previous work [1]. NTPs and TIPP are included in the IVT reaction but are not shown in the schematic in **A**. Raw fluorescence values were standardized to µM FITC.

To explore whether coacervates are compatible with the more complex transcriptional regulation process that underlies ROSALIND, we focused on the aTF TetR, which unbinds from the operator *tetO* in the presence of either ligands anhydrotetracycline (aTc) or tetracycline (TCY) [72]. To test whether TetR repression of transcription is functional in the presence of coacervates, we titrated a range of TetR concentrations into a room-temperature IVT reaction transcribing 3WJdB downstream of a *tetO* operator sequence. We observed that TetR can successfully repress transcription of 3WJdB, albeit at higher TetR concentrations when coacervates are added (**Supplementary Fig. 11A-B**), potentially due to coacervate-mediated amplification of RNAP activity that has escaped repression.

We next tested whether ligand-mediated derepression is functional with coacervates. After adding aTc to the IVT repressed by 150 nM TetR dimer, kinetic traces demonstrated that aTc can induce 3WJdB production at room temperature with coacervates. Excitingly, coacervates also accelerate the signal kinetics of aTc biosensing, consistent with our constitutive IVT experiments, reducing the time to visible signal (1 μM FITC) from 117 to 54 mins and resulting in 2.4-fold increased signal after 2 hours (**Fig. 4B, Fig. 4H-I** and **Supplementary Fig. 12**). We repeated these experiments, inducing TetR depression with TCY, and achieved a time to visible signal reduction from 146 to 64 mins, and 2.5X enhancement of signal at 2 hours (**Fig. 4C** and **Fig. 4H-I**). We additionally confirmed that coacervates do not alter the specificity of TetR biosensing (**Supplementary Fig. 13**).

Previous work showed that at 37 °C, ROSALIND kinetics can be enhanced by switching reporters from 3WJdB to a toehold-mediated strand displacement (TMSD) readout, which involves RNA-mediated strand displacement of a DNA duplex [19]. To test how coacervates perform with respect to TMSD, we compared our results to TMSD-mediated aTc-sensing reactions and found that TMSD outputs are slightly slower than 3WJdB at room temperature, and that reactions with 3WJdB and coacervates significantly outperform those with TMSD at room temperature (**Supplementary Fig. 14**).

We next determined whether coacervate-mediated enhancement is compatible with different families of aTFs that detect various classes of ligands. In addition to *E. coli* TetR, we chose *S. elongatus* SmtB, *B. subtilis* CsoR, *Comanomas spp.* MobR, and *P. putida* TtgR. These aTFs span four distinct families of transcriptional regulators (TetR [73], ArsR/SmtB [74], CsoR/RcnR [75], and MarR [76]) and sense zinc, copper, 3-hydroxybenzoic acid, and naringenin, respectively. We placed the cognate operator sequence of each aTF after the T7 RNAP promoter and immediately before the DNA encoding for 3WJdB. We found that coacervates enhance the unregulated transcription of all tested DNA templates. To boost the enhancement effect observed for the template containing the *csoO* operator, we also increased the DNA template concentration. (**Supplementary Fig. 11A**). Next, we titrated each of the aTFs into IVT reactions and established repression for each aTF-operator pair at 6 μM SmtB dimer, 4.5 μM CsoR tetramer, 32 μM MobR dimer and 10 μM TtgR dimer, with coacervate-mediated differences in repression similar to our findings with TetR (**Supplementary Fig. 11C-F**). We then demonstrated that ROSALIND with coacervates can be used to sense zinc, copper, 3-hydroxybenzoic acid, and naringenin, with accelerated room temperature reaction rates (**Fig. 4D-G**) that reduce time to visible signal by an average of 68 minutes (**Fig. 4H**) and result in an approximately 2X enhancement in signal magnitude after 2 hours (**Fig. 4I**).

Together, these results demonstrate that spermine-PAA coacervates are compatible with regulated IVT biosensing that leverages multiple transcription factor families and can accelerate the detection of diverse ligands.

### Coacervates can increase transcriptional biosensor sensitivity

Many small molecules are selectively enriched within biological phase-separated systems [37, 77], with recent research suggesting that hydrophobicity-related features are among the strongest predictors of small-molecule partitioning [78]. Based on these observations, we hypothesized that incorporating coacervates into transcriptional biosensor reactions could alter detection sensitivity by selectively concentrating ligands, with the trend and magnitude dependent on ligand hydrophobicity. To investigate this hypothesis, we generated room-temperature dose-response curves for aTc and TCY, which share the transcription factor TetR and the DNA template but differ in ligand hydrophobicity. Using 150 nM TetR, we observed that the EC50 of the dose-response curve, defined as the ligand concentration that generates half maximal signal, for the hydrophobic molecule aTc (**Fig. 5A**) was slightly increased, while the EC50 of that for the more polar molecular TCY (**Fig. 5B**) was reduced by 1.3-fold. This finding confirms that coacervates alter biosensor sensitivity in response to ligand characteristics. To investigate how coacervates impact the biosensor sensitivity to other ligands, we also generated room-temperature dose-response curves for zinc, copper, 3-hydroxybenzoic acid, and naringenin. We observed that the EC50 values of the dose-response curves decreased by 1.4-, 1.1-, 1.1-, and 1.5-fold, respectively, when coacervates were added to biosensor reactions run at room temperature (**Fig. 5C-F**). Notably, coacervates increased the maximal signal observed for all dose-responses.

**Figure 5:**
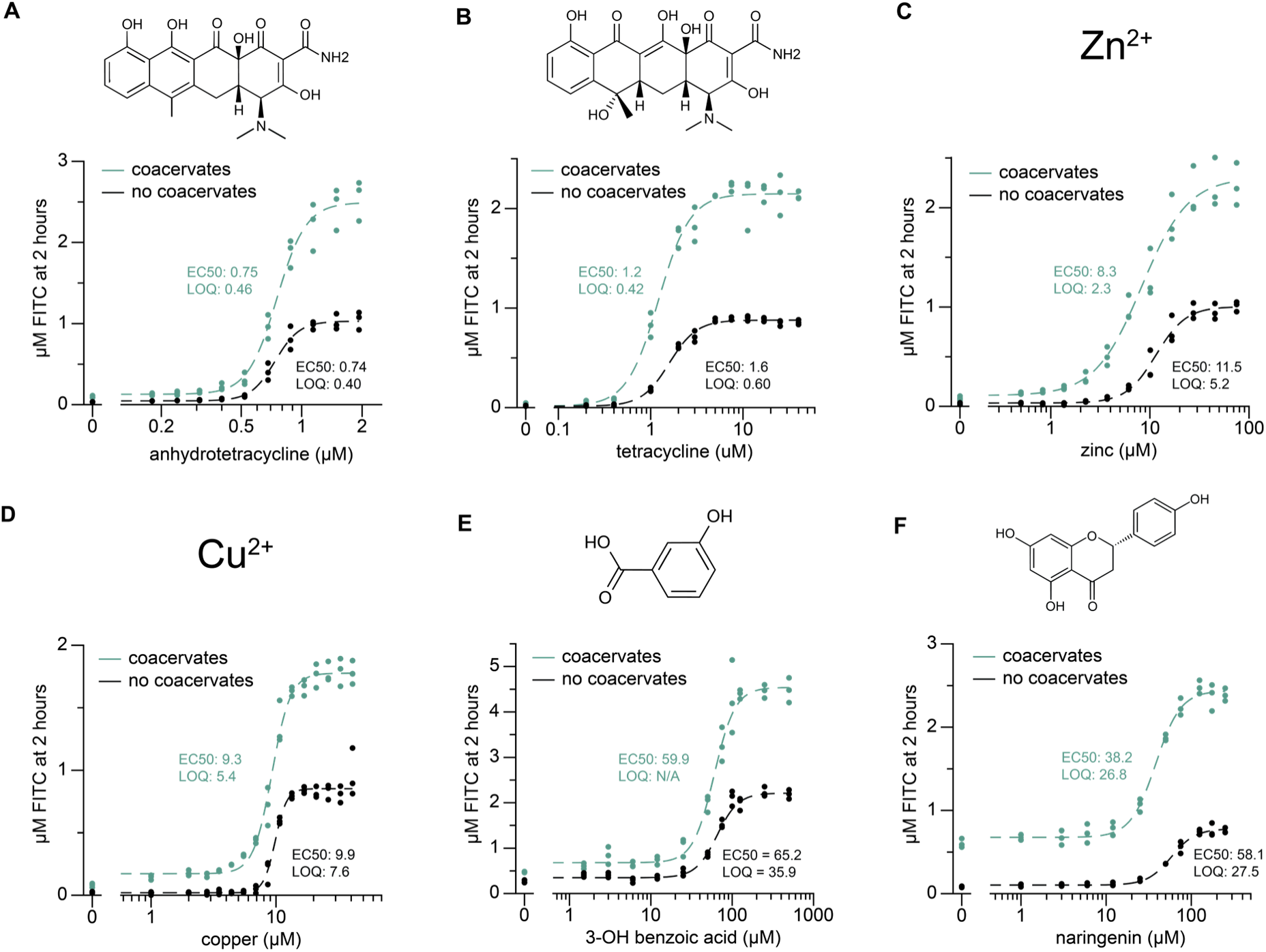
Coacervates can enhance cell-free biosensor sensitivity. **A,B**, Coacervates influence the sensitivity of ligand detection based on ligand hydrophobicity. The dose-response EC50 for the less polar molecule anhydrotetracycline slightly increases from 0.74 to 0.75 (**A**), whereas the EC50 for the more polar molecule tetracycline decreases from 1.6 to 1.2 (1.3-fold) (**B**). The limits of quantification (LOQs) follow the same trend as the EC50 values. Reactions in **A,B** contained the same DNA template and TetR concentration. **C-F**, Dose-responses with zinc (**C**), copper (**D**), 3-hydroxybenzoic acid (**E**), and naringenin (**F**) demonstrate coacervate-mediated reductions in EC50 of 1.1 to 1.5-fold, with similar reductions in LOQ. The EC50 is defined as the analyte concentration at which half-maximal signal is achieved. The limit of quantification (LOQ) is defined as the analyte concentration corresponding to ten standard deviations above the blank mean. N/A indicates where the LOQ could not be determined (see **Supplementary Data 3** and **Supplementary Data 4**). Both EC50s and LOQs were calculated using Hill equation fits, which are illustrated by the dashed curves (**Methods**). Reactions with coacervates show increased maximal signal for the detection of all ligands. All reactions were run at room temperature (∼25 °C), with the data shown collected after 2 hours of incubation. DNA template and aTF concentrations used in each reaction are listed in **Supplementary Table 3**, and the DNA and protein sequences are listed in **Supplementary Data 1**. The data shown in **A-F** are *n* = 3 independent biological replicates, each plotted as a point with raw fluorescence standardized to µM FITC.

In previous work, we showed that transcriptional biosensors can be sensitized by reducing the aTF concentration while keeping the aTF-to-DNA ratio constant, which requires signal amplification to generate sufficient signal at reduced DNA template concentrations [5]. Because coacervates amplify IVT signal, we sought to apply this approach as a second, independent strategy leveraging coacervates to sensitize detection. Maintaining the aTF to DNA ratio to around 250, we discovered that an IVT reaction with coacervates, 2.5 µM smtB dimer and 10 nM DNA can achieve the same signal as a control reaction with 25 nM DNA template (**Supplementary Fig. 15**). This system yields a 3.7-fold improvement in EC50 compared to a control reaction containing 25 nM DNA template and 6 µM smtB (**Supplementary Fig. 15A**). Because IVT without coacervates can be repressed with less aTF, we also benchmarked to a control IVT with 25 nM DNA template and 2 µM smtB, which is sufficient to represses a control IVT reaction (**Supplementary Fig. 11C**), and we observed a 1.8-fold improvement in EC50 with this comparison (**Supplementary Fig. 15B**). These findings suggest that coacervate-mediated amplification can be repurposed to sensitize detection.

EC50 provides a functionally useful measure of sensitivity, particularly in the absence of instrumentation, reflecting the concentration required to elicit a substantial response across the dynamic range. However, because EC50 is defined as the concentration yielding 50% of the maximal response, higher signal amplitudes impose higher signal thresholds for EC50 determination. Therefore, we also assessed biosensor sensitivity using the limit of detection (LOD) and limit of quantification (LOQ) metrics (**Figure 5, Supplementary Fig. 16** and **Supplementary Data 3**). The trend in sensitization was broadly consistent across EC50, LOD, and LOQ, although the magnitude of the fold change varied, likely because LOD and LOQ are statistical tests that quantify small differences near the background. As a direct, field-relevant measure of sensitivity, we also determined the analyte concentration required to induce visible signal in 2 hours at room temperature and demonstrated that this is reduced with coacervates for all analytes, noting that the fold change magnitudes that could be calculated are limited, as many dose-responses without coacervates fail to reach visible signal in 2 hours at any analyte concentration (**Supplementary Fig. 16** and **Supplementary Data 3**). Together, these results demonstrate that, in addition to accelerating reactions, coacervates can enhance the sensitivity of analyte detection.

### Coacervate-mediated enhancement of biosensing is robust in point-of-use formats

Freeze-drying of cell-free biosensors enables cold-chain-free storage, facilitating distribution and improving accessibility for use in field settings [17, 18, 79, 80]. After freeze-drying, water-contaminant biosensors can, for example, be shipped and rehydrated with a drop of water at the point of use [11, 13]. Unfortunately, freeze-drying can both slow reaction kinetics and decrease endpoint signals [5, 11, 14, 81], and both these consequences can be exacerbated by matrix effects from real-world samples [1, 5, 19]. Therefore, we explored whether coacervates could be applied to enhance signal in these more challenging conditions and leveraged to mitigate their effects.

To investigate this hypothesis, we assembled IVT reactions with or without coacervates and freeze-dried them as described previously [1]. Importantly, all components of the reaction were freeze-dried together, enabling a ‘single-pot’ format that was subsequently rehydrated with 20 µL of water and incubated at room temperature (**Fig. 6A**). We observed that rehydration of freeze-dried unregulated IVT reactions with coacervates demonstrated improved kinetics compared to the no-coacervate control (**Supplementary Fig. 17**). We then freeze-dried the zinc-sensing ROSALIND biosensor with coacervates, with an increased smtB concentration to ensure robust repression after lyophilization and rehydration in sample matrices (**Supplementary Table 3**). We demonstrated that the kinetic speedup is maintained after rehydration with both lab-grade water and unprocessed Lake Michigan water spiked with zinc (**Fig. 6B** and **Fig. 6E**), with the time to visible signal reduced by 1.4 and 3.5 hours, respectively (**Fig. 6C** and **Fig. 6F**). Finally, we performed a dose-response of freeze-dried zinc-sensing ROSALIND reactions rehydrated with various amounts of lab or unprocessed Lake Michigan water and showed that the sensitivity enhancement is maintained (**Fig. 6D** and **Fig. 6G**).

**Figure 6:**
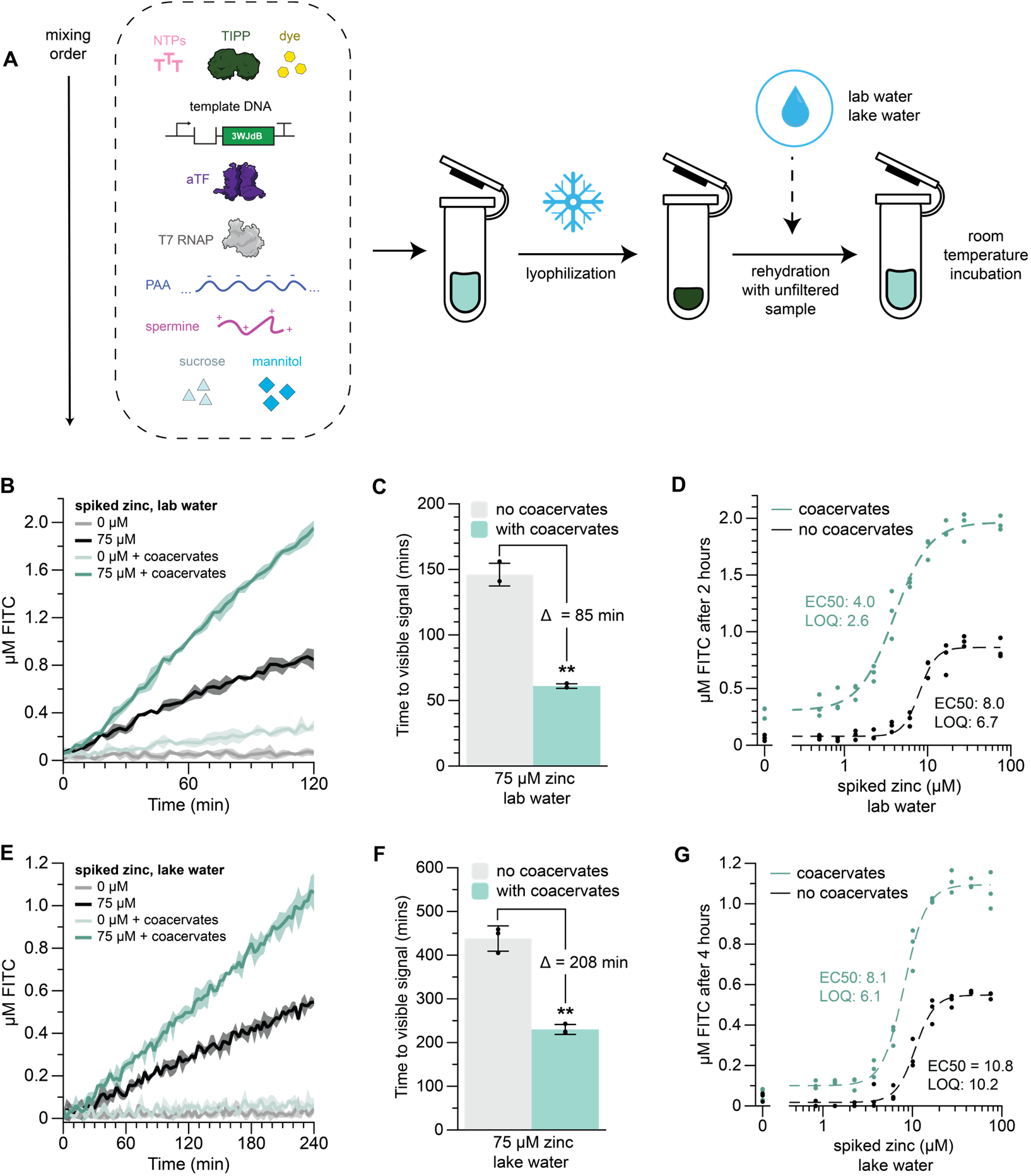
Coacervates accelerate and sensitize target detection in real-world matrices. **A**, ROSALIND reactions with spermine-PAA coacervates were freeze-dried, rehydrated with either lab water or unfiltered Lake Michigan water (spiked with zinc), and then incubated at room temperature. **B-D**, Coacervate-mediated enhancement of biosensing is maintained after lyophilization. Zinc-sensing reactions with and without coacervates were lyophilized and rehydrated with 20 µL of laboratory water spiked with zinc. Reactions with coacervates demonstrated an 85-min reduction in time to visible signal (**B,C**), and a 2-fold decrease in EC50 (**D**). **E-G**, Coacervates accelerate and sensitize zinc biosensing in spiked Lake Michigan water. Reactions with coacervates demonstrated a 3.5-hour reduction in time to visible signal (**E,F**), and a 1.3-fold decrease in EC50 (**G**). DNA template and aTF concentrations used in each reaction are listed in **Supplementary Table 3**, and the DNA and protein sequences are listed in **Supplementary Data 1**. All reactions were run at room temperature (∼25 °C). For the data in **B,E**, the shading indicates the average value of n=3 independent biological replicates +/- standard deviation. The data shown in **C,F** summarize the kinetic traces shown in **B,E**, with bar heights representing the average of n=3 replicates, each plotted as a point, and error bars indicating the standard deviation. A two-tailed, heteroscedastic Student’s *t*-test was used to compare the time to signal between conditions with and without coacervates. The *P* value range in **C,F** is indicated by asterisks (\*\*\**P* < 0.001, \*\**P* = 0.001–0.01, \**P* = 0.01–0.05, ns otherwise). Time to visible signal is defined as the time to 1 µM FITC, based on 3WJdB fluorescence visualized in previous work [1]. The data shown in **D,G** are n=3 independent biological replicates, each plotted as a point with raw fluorescence standardized to µM FITC. The EC50 and LOQ values were calculated using the Hill equation fits, which are illustrated by the dashed curves (**Methods**).

Unlike previous ROSALIND studies [1, 5, 19], we performed all experiments at room temperature, demonstrating that coacervates yield enhancements without requiring a field-portable incubator. Together, these results show that coacervate-mediated IVT enhancement is robust to freeze-drying and could be easily interfaced with biosensors to enhance the speed and sensitivity of contaminant detection at the point-of-use.

## DISCUSSION

In this study, we demonstrate that coacervation can be interfaced with IVT to accelerate and sensitize cell-free biosensing. To induce phase separation, IVT reactions were mixed with polyelectrolytes spermine and PAA. Room-temperature IVT reactions with coacervates demonstrated a 2-fold increase in the output signal, achieving fluorescence values greater than those of controls incubated at 37 °C (**Fig. 1I**). We show that coacervates are compatible with several aTF families to sense a wide range of compounds, with an average biosensor time-to-signal reduction of 68 minutes and a 1.2-fold reduction in EC50 (**Fig. 4** and **Fig. 5**). Coacervates were also amenable to single-pot lyophilization with biosensor components, and reduced time-to-signal by 3.5 hours when challenged with real-world matrices (**Fig. 6F**).

To our knowledge, this is the first phase-separated system that leverages the self-assembly of polyelectrolytes to enhance *in vitro* transcription. While transcriptional and translational kinetics are often improved when phase separation is induced within cells [32, 33, 49], coacervation *in vitro* tends to inhibit these same processes [46, 48, 49]. Consistent with prior reports [48, 55], our work suggests that polycation poisoning of *in vitro* systems is a contributing factor (**Fig. 1E**). Interestingly, the polycation we use, spermine, is an endogenous polyamine that improves RNA synthesis in reconstituted eukaryotic condensates [82] and has been successfully applied to generate coacervates in other *in vitro* contexts [39, 59]. Enhanced polymer biocompatibility may thus explain this system’s ability to enhance IVT, although we importantly show that adding spermine alone cannot account for the transcriptional improvements we describe (**Fig. 1F, G**).

Interestingly, our coacervate system may serve as a model for more complex phase separation phenomena in biology. Specifically, we identify that coacervation in our system is induced by transcription via NTP hydrolysis (**Fig. 2**) and in turn accelerates transcription through co-concentration of DNA and RNAP (**Fig. 3**). This process echoes recent findings showing that native transcription regulates phase separation in cells [82–84], which in turn modulates transcription through recruitment of chromatin [85] and RNA polymerase [86]. We anticipate that the unique mechanism associated with our coacervate system may enhance its relevance to the study of transcriptional condensates, potentially serving as a simple model for testing biophysical theories relevant to dynamic cellular condensation.

A potential limitation of our approach of applying coacervates to biosensing applications is their susceptibility to external conditions. Coacervation can be sensitive to temperature, salt and pH [87–89], and may fail to occur in complex sample matrices. However, coacervates are currently in development to withstand unfavorable environments, including human blood [90] and the gastrointestinal tract [91]. We also achieve meaningful coacervate-mediated improvements when testing with real-world lake water samples (**Fig. 6E-G**). We expect that recent advances in membrane encapsulation of coacervates could also be applied to improve coacervate robustness and utility for biosensing [92, 93]. Furthermore, we note that when transcription is weakened, coacervation in our system is delayed and the enhancement effect is reduced, but we can bolster the basal transcription by tuning the DNA or RNAP concentration (**Fig. 2H-I**).

Key features of our approach include simplicity and cost. Both spermine and PAA are inexpensive, commercially available polymers, and their inclusion in IVT reactions to form coacervates requires no additional steps beyond mixing. By leveraging self-assembly to improve sensing, coacervates also offer significant cost advantages compared with other enhancement approaches to cell-free biosensing, including CRISPR-Cas technologies, which require additional purified enzymes [24], or TMSD, which requires expensive modified oligonucleotides [19]. We show that IVT reactions with coacervates significantly outperform TMSD at room temperature, although TMSD can enable additional functionalities beyond kinetic improvement (**Supplementary Fig. 14**).

We also observed small improvements in detection sensitivity for most analytes detected in this study (**Fig. 5**). Ligand sensitivity is likely driven by both ligand and aTF partitioning, as well as by crowding effects, and is likely to vary substantially across target molecules. Recent work suggests that partitioning can span a millionfold range across different small molecules, yet can be predicted relatively independently of coacervate chemistry [78]. Additional work investigating the sensing of other ligands will be necessary to fully understand the impact of coacervation on sensitivity, and could leverage biosensor schemes that apply promiscuous aTFs that sense a variety of ligands to control for aTF partitioning. Nevertheless, the data here demonstrates that coacervates can improve assay sensitivity for a variety of ligands detected by many different aTF families. We have previously shown that schemes that enhance fluorescence output can be reconfigured to improve sensitivity [5], and we also successfully leverage this strategy to repurpose the signal enhancement provided by coacervates to enhance the sensitivity of zinc sensing (**Supplementary Fig. 15**).

We believe that this work opens the door to leveraging phase separation to improve other *in vitro* cell-free technologies. Specifically, coacervates could be combined with cell-free biosensing reactions that leverage DNA nanotechnology-encoded logic gates or amplification circuits to further increase speed, improve sensitivity, or enable computation. In addition, coacervates could enable multiplexed biosensing, as recent demonstrations have shown that specific DNA templates can be anchored to distinct phase-separated droplets [52]. We also anticipate interesting applications of spermine-PAA coacervates as model compartments to study the formation and dynamics of biological condensates. For example, stimulus-responsive *in vitro* transcription could be coupled with processes such as DNA replication, recently demonstrated in coacervates [94], to provide a simplified model for the concurrent replication and transcription in the nucleolus [95]. Partitioning of key molecules in this system could then be investigated with recent advances in label-free and NMR methods [96, 97]. We believe these future studies could yield interesting insights into biological phase separation, defining fundamental mechanisms that could extend to research on primitive protocells and the origins of life.

Taken together, this work establishes an interface between self-assembling coacervate materials and modular, *in vitro,* cell-free biosensing. We anticipate that the advantages of this approach, including affordability, simplicity, enhancements in kinetic performance and sensitivity, and compatibility with reduced equipment, room-temperature operation, modular aTFs, and lyophilization, will make coacervation-enhanced transcriptional technologies a promising tool for applications in both biology and biotechnology.

## METHODS

### Materials

Spermine (≥97%) was purchased from Sigma-Aldrich. Spermidine trihydrochloride (≥99%) was purchased from Thermo Scientific. Poly(diallyldimethylammonium chloride) (PDADMAC, 8500 g/mol) was purchased from Polysciences, Inc. Poly(acrylic acid sodium salt) (PAA, 5100 g/mol) was purchased from Millipore Sigma. Poly(ethylene glycol) (PEG, 8000 g/mol) was purchased from Sigma-Aldrich. Charge density was calculated by assuming that all the protonable nitrogen atoms in the polymers are protonated at the reaction pH. T7 RNA polymerase (RNAP) and thermostable inorganic pyrophosphatase (TIPP) was purchased from New England Biolabs. Tris-buffered nucleotide triphosphates (NTPs, pH 7.3-7.5) were purchased from ThermoScientific. DFHBI-1T was purchased from Tocris Bioscience. UltraPure DNase/RNase-Free distilled water was purchased from Invitrogen. Sucrose and D-mannitol were purchased from Sigma-Aldrich.

### Strains and growth medium

The BL21(DE3) competent *E. coli* cell strain was used for recombinant protein expression. The K12 *E. coli* cell strain (NEB Turbo Competent *E. coli*, New England Biolabs) was used to propagate plasmids encoding transcription templates. Luria broth supplemented with the appropriate antibiotic(s) (100 µg mL^−1^ carbenicillin, 100 µg mL^−1^ kanamycin, and/or 34 µg mL^−1^ chloramphenicol) was used as the growth medium, unless noted otherwise.

### Plasmids and genetic parts assembly

Protein expression plasmids to produce TetR, SmtB, CsoR, MobR and TtgR were cloned previously and were designed to overexpress recombinant proteins as N-terminus or C-terminus His-tagged fusions [1]. Protein expression plasmids for SmtB, CsoR and MobR have a tobacco etch virus (TEV) protease recognition sequence between the aTF and the 6X His-tags.

Plasmids encoding the transcription templates were also cloned previously [1], and were designed to include a T7 RNAP promoter, an aTF operator site, the three-way junction dimeric Broccoli (3WJdB) coding sequence, and a T7 terminator (T7-operator-3WJdB-T). Transcription templates were generated by PCR amplification (Phusion High-Fidelity PCR Kit, New England Biolabs) from the plasmids to include a 5′-region upstream of the T7 RNAP promoter and a 3′-region ending with the T7 terminator, using the DNA primers (Integrated DNA Technologies) listed in **Supplementary Data 1**. The PCR-amplified templates were purified (QIAquick PCR purification kit, Qiagen) and verified for the presence of a DNA band of expected size on a 2% Tris–acetate–EDTA–agarose gel. The concentrations of all DNA templates were determined using a NanoDrop One Spectrophotometer (ThermoScientific). All plasmid sequences were verified using Nanopore sequencing (Quintara Biosciences) prior to use.

All plasmids and DNA templates were stored at 4 °C until use. A spreadsheet listing the sequences and the Addgene accession numbers of all oligos and plasmids used in this study, respectively, is provided in **Supplementary Data 1**.

### Replicates

Where replicates are shown, they represent independent biological replicates (n = 3), defined as reactions assembled from separate master mixes and aliquoted into distinct tubes. Replicates were processed and analyzed separately (e.g., in separate wells or measurements). Lyophilized replicates were lyophilized in the same chamber but rehydrated separately.

### *In vitro* transcription and ROSALIND reactions

*In vitro* transcription (IVT) reactions were set up by adding the following components listed at their final concentrations: transcription buffer (**Supplementary Table 2**), 0.2 mM DFHBI-1T, 11.4 mM Tris-buffered NTPs, 0.3 units (U) of TIPP, 25 nM DNA transcription template (unless indicated otherwise), and UltraPure H_2_O to a total volume of 20 µL. ROSALIND reactions additionally included a purified aTF at the indicated concentration (**Supplementary Table 3**). All components listed above were first assembled and preincubated at 37 °C for 15 min. Immediately before plate reader measurements, 10 U μL^−1^ (unless noted otherwise) of T7 RNAP (New England Biolabs) and, optionally, a ligand at the indicated concentration (**Supplementary Table 3**) were added to the reaction. For IVT and ROSALIND reactions with coacervates, polyelectrolytes were added after addition of T7 RNAP, but before addition of any ligand, at the concentrations specified in **Supplementary Table 1**. For IVT reactions containing PEG, PEG was added to replace at equal mass either PAA alone (16.2 μg) or PAA with spermine (53.8 μg) after the addition of T7 RNAP. All room temperature reactions were run at ambient laboratory temperature, which ranged between 24 and 26 °C; the exact temperature corresponding to each room-temperature experiment is provided in the **Source Data**. The reactions were characterized on a plate reader as described in the “Plate reader quantification and fluorescence standardization” section. See **Supplementary Data 2** for a Microsoft Excel worksheet describing the assembly of a ROSALIND reaction with spermine-PAA coacervates.

### Coacervate formation studies

To understand how IVT reaction components may inhibit immediate coacervate formation, IVT reactions with coacervates were assembled as previously described, without T7 RNAP where indicated to prevent transcription. Components of the IVT mixture were omitted one at a time or titrated as indicated, and the turbidity (absorbance at 600 nm) of the assembled mixtures was immediately measured after mixing using a plate reader at room temperature. Coacervate dispersions were also assembled by mixing transcription buffer, 20 mM charge concentration PAA, and 4 mM spermine. To characterize the formation of coacervates within active IVT reactions, transcription reactions with and without coacervates were prepared as described above, with 10 U μL^−1^ (1X) or 20 U μL^−1^ (2X) of T7 RNAP, and incubated inside a plate reader at either room temperature or 37 °C. Turbidity (absorbance at 600 nm) was measured at 3-min intervals for 4 hours, with plates shaken for 10 seconds prior to each measurement. Samples were imaged using the 40X air objective on a Nikon Ti2 Eclipse epifluorescence microscope to obtain DIC images.

### Purified RNA studies

Purified RNA was generated through IVT using the DNA template T7-tetO-3WJdB-T, with the following components: transcription buffer (**Supplementary Table 2**), 11.4 mM Tris-buffered NTPs, 1.5 U TIPP, 50 nM transcription template, 10 U μL^−1^ of T7 RNAP, and water to a total volume of 100 μL. Three IVT reactions were incubated at 37 °C for four hours, ethanol-precipitated, and gel-purified by resolving on a 7% urea–PAGE–TBE gel, isolating the major product band (expected size 223 nucleotides), and eluting at 4 °C overnight in UltraPure H_2_O. The eluted RNA was then ethanol precipitated, resuspended in UltraPure H_2_O, quantified using the Qubit RNA BR Assay Kit (Invitrogen), and stored at 4 °C for use. The RNA sequence is available in **Supplementary Data 1**.

To investigate whether RNA generation influences coacervate formation, purified RNA was added to IVT reactions without T7 RNAP and assembled with NTPs at 11.4 mM (1X) or 5.7 mM (0.5X). Turbidity (absorbance at 600 nm) was measured immediately after assembly using a plate reader. To characterize the effect of spermine-PAA coacervates on purified 3WJdB fluorescence, purified RNA was added to DFHBI-1T, with or without coacervates, in mixtures totaling 20 μL. For mixtures not containing coacervates, mixture components were assembled in the following order: transcription buffer (**Supplementary Table 2**), 0.2 mM DFHBI-1T, purified tetO-3WJdB-T RNA, and UltraPure water. For mixtures containing coacervates, reaction components were assembled in the following order: transcription buffer (**Supplementary Table 2**), 0.2 mM DFHBI, purified tetO-3WJdB-T RNA, water, 20 mM charge concentration PAA, and 4 mM spermine. 19 μL of each sample was then loaded into a 384-well plate, and the fluorescence of the mixture after 2 hours was measured as described in the “Plate reader quantification and fluorescence standardization” section.

### RNA quantification from IVT reactions

For the RNA products quantified in **Fig. 3A** and **Supplementary Fig. 4**, 20 μL IVT reactions with and without coacervates were set up as described above, except DFHBI-1T was replaced with DMSO. All IVT reactions contained 25 nM of the T7-tetO-3WJdB-T DNA template (**Supplementary Data 1**). After 2 hours at room temperature, transcription was stopped by adding 80 μL of TRIzol Reagent (Invitrogen). After adding 16 μL chloroform (Acros Organics), samples were vortexed, incubated at room temperature for 10 mins, and centrifuged for 15 mins at 4 °C, after which 50 μL of the aqueous phase was removed. Samples were then ethanol precipitated with 1 μL glycogen coprecipitant (GlycoBlue Blue Coprecipitant, Invitrogen), resuspended in 1× TURBO DNase buffer (Invitrogen) containing 2 μL TURBO DNase (Invitrogen), and incubated at 37 °C for 2 hours. Following DNA digestion, RNA extraction and purification was performed using 200 μL TRIzol Reagent and 40 μL chloroform, and 120 μL of the aqueous phase was extracted.

Samples were then ethanol precipitated with 1 μL glycogen coprecipitant and resuspended in 10 μL of UltraPure water.

The RNA concentration of each sample was quantified using a Qubit RNA Broad Range Assay Kit (ThermoFisher Scientific) following the manufacturer’s instructions. To investigate sample differences via gel electrophoresis, 0.7 µL of each sample was mixed with water and 2X RNA loading dye (New England Biolabs), added to a 7% urea–PAGE–TBE gel, and run alongside a Century-Plus RNA ladder (Invitrogen). Gels were stained with SYBR Gold Nucleic Acid Gel Stain (Invitrogen) and imaged using a ChemiDoc Touch Gel Imaging System (Bio-Rad Image Lab Touch software v.1.2.0.12). Nucleic acid band intensity was quantified by defining a region of interest (ROI) sized to the largest band, centering the ROI on each band, and measuring the mean grey value in greyscale images using ImageJ/Fiji [98]. The intensity value for each lane was corrected for background signal by subtracting the mean gray value of an empty region below each band.

### DNA sequestration studies

To characterize DNA sequestration, we adapted methods from the literature to assay molecular sequestration within coacervates [37, 48]. 100 µL mixtures of either (1) 25 nM DNA, (2) 25 nM DNA with spermine-PAA coacervates, or (3) spermine-PAA coacervates, were prepared in transcription buffer containing the DNA sequence and coacervate composition as described in **Supplementary Data 1** and **Supplementary Table 1**, respectively. The order of mixing was transcription buffer, DNA, water, PAA, and spermine. Each mixture was allowed to equilibrate for 3 mins at room temperature, after which 4 µL of each mixture was collected as a pre-centrifugation sample. Mixtures were then centrifuged for 5 min at 5000 RCF to pellet the droplets, and 4 µL was isolated from the supernatant. The six samples (three collected prior to centrifugation, and three collected from the supernatant after centrifugation) were then diluted in water and 6X Purple Gel Loading Dye without SDS (New England Biolabs) into a 2% Tris–acetate–EDTA–agarose gel cast with GelRed (Biotium). After running, gels were imaged using a ChemiDoc Touch Gel Imaging System. DNA band intensity was quantified in the same manner described for RNA bands.

To confirm that the centrifugation process completely removes droplets, mixtures were prepared as described above, with 20 µL of each mixture isolated before centrifugation and 20 µL isolated from the supernatant after centrifugation. Samples were then placed into a 364-well plate for turbidity quantification (absorbance at 600 nm). To estimate the coacervate dense phase volume after centrifugation, volume standards were prepared by placing 0.5 µL, 1 µL, 2 µL, and 3 µL solutions of 0.024% bromophenol blue (Sigma-Aldrich) in formamide (Invitrogen) into tubes and briefly centrifuging to collect the drops at the bottom of the tubes. The volume of the coacervate after centrifugation was then visually compared to the volume standards. All images were taken on white printer paper inside a Glendan Portable Photo Studio Light Box, and experiments were performed in triplicate.

### Fluorescence microscopy

To investigate the localization of IVT components, we prepared coacervate mixtures containing fluorescently-labeled DNA or T7 RNAP. Fluorescent DNA was prepared through PCR amplification (Phusion High-Fidelity PCR Kit, New England Biolabs) with a forward DNA primer containing a 5’ Alexa Fluor 546 modification (Integrated DNA Technologies), with the sequence as described in **Supplementary Data 1**. Fluorescent T7 RNAP was prepared by expressing and purifying T7 RNAP (**Supplementary Methods**) and conjugating the protein to DyLight 405 using NHS ester-amine chemistry (DyLight Antibody Labeling Kits, Thermo Scientific). 40 µL samples for imaging were prepared by sequentially combining transcription buffer, labeled DNA and/or labeled T7 RNAP, water, 20 mM charge concentration PAA, and 4 mM spermine, as described in **Supplementary Table 4**. After assembly, the tube was briefly shaken for approximately 2 s, and the entire sample was transferred by pipette into a single well of a chambered slide (Ibidi µ-Slide 18 Well). A coverslip was placed over the chamber to minimize evaporation during imaging. Prior to use, the slide was treated to impart hydrophobicity according to an established protocol [99].

Fluorescence microscopy experiments were performed using an SP8 resonant scanning confocal microscope (Leica Microsystems) and imaged with a 40× oil-immersion objective (HC PL APO CS2 40x, 1.3 NA, oil-immersion, Leica Microsystems). Samples were illuminated with a tunable white-light laser set to 405 nm to excite the DyLight 405 dye or 557 nm to excite the Alexa Fluor 546 dye. Photons were detected with a PMT detector and filtered with 410–564 grating (DyLight 405 dye) or 562–764 nm grating (Alexa Fluor 546 dye). Single-frame images were taken either in a 320 x 320 or 672 x 672 pixel array, corresponding to image areas of 18.5 µm^2^ or 47.4 µm^2^, respectively. Brightfield images were collected in tandem with fluorescence imaging using a transmitted light detector (TLD). ImageJ was used to apply false-color mapping to the raw grayscale images using cyan and yellow color schemes. When the Dylight 405 and AlexaFluor 546 imaging agents were used, the display settings were changed to min = 0 and max = 50 (cyan) and min = 0 and max = 120 (yellow), respectively. These display settings are used in **Fig. 3C, Supplementary Fig. 8B**, and **Supplementary Fig. 9B**. Merged channel images were generated in ImageJ using the same previous display settings for each color channel.

### Expression and purification of T7 RNAP and aTFs

T7 RNAP and aTFs were expressed and purified as previously described [1, 100]. Briefly, sequence-verified protein expression plasmids were transformed into BL21(DE3) competent *E. coli* cells. 1-4 L cell cultures were grown at 37 °C, induced with 0.5-1 mM of isopropyl-β-d-thiogalactoside (IPTG) at an optical density (600 nm) ∼ 0.5, and grown for four additional hours at 37 °C. The cultures were then pelleted by centrifugation, flash-frozen in liquid nitrogen, and stored at −70 °C. On the day of purification, pellets were thawed and resuspended in a lysis buffer containing a protease inhibitor, then lysed by ultrasonication or homogenization, and insoluble material was removed by centrifugation. Clarified supernatants were then purified as described in **Supplementary Table 5**. His-tags were cleaved for the CsoR protein to avoid polyhistidine-copper binding, but retained for other aTFs to enhance purification yields. For aTFs, eluted fractions were concentrated and buffer exchanged using centrifugal filtration (Amicon Ultra Centrifugal Filters, Milipore Sigma). Protein concentrations were determined using the Qubit Protein Assay Kit (Invitrogen). The purity and size of the proteins were validated by SDS–PAGE and purified proteins were stored at −20 °C. Purified T7 RNAP was subsequently labeled with a fluorophore and used solely for fluorescent imaging. A list of protein sequences is provided in **Supplementary Data 1**, and detailed purification protocols are provided in the **Supplementary Methods**.

### Lyophilization

Before lyophilization, the caps of PCR tubes were punctured with a pin to create three holes. Lyophilization of ROSALIND reactions without coacervates was performed as described previously [1], by assembling all the components of IVT with the appropriate aTF, with 50 mM sucrose and 250 mM D-mannitol as lyoprotectants. Lyophilization of ROSALIND reactions with coacervates additionally included 4 mM spermine and 20 mM charge concentration PAA, added before the 50 mM sucrose and 250 mM D-mannitol. The assembled reaction tubes were transferred into an aluminum block and wrapped in aluminum foil. The block was then submerged in liquid nitrogen and transferred to a FreeZone 2.5-L BenchTop Freeze Dry System (Labconco) for overnight freeze-drying at a condenser temperature of −85 °C and a pressure of 0.04 mbar. When rehydrating the lyophilized reactions, 20 µL of water (optionally containing a ligand) was added to the tube to resuspend the pellet. The reactions were then characterized on a plate reader as described in the “Plate reader quantification and fluorescence standardization” section.

### Lake water sample collection

Lake water samples were collected in Evanston, IL, by submerging a 50 mL Falcon tube into Lake Michigan. Samples were capped, brought back to the lab, and stored at room temperature until use.

### Plate reader quantification and fluorescence standardization

A National Institute of Standards and Technology-traceable standard (Invitrogen) was used to convert arbitrary fluorescence measurements to micromolar-equivalent fluorescein (µM FITC). Serial dilutions from a 52 µM stock were prepared in 100 mM sodium borate buffer at pH 9.5, including a 100 mM sodium borate buffer blank (total of 6 samples). For each concentration, three replicates were created, and the fluorescence values were read at an excitation wavelength of 472 nm and emission wavelength of 507 nm for 3WJdB-activated fluorescence, or an excitation wavelength of 495 nm and emission wavelength of 520 nm for 6-FAM TMSD outputs (Synergy H1, BioTek Gen5 v.3.10).

The fluorescence values from the three replicates were averaged for each fluorescein concentration, and a linear regression was performed in Datagraph 5.2 to correlate the average fluorescence values (arbitrary units) with fluorescein concentration. For each plate reader, excitation, emission, and gain setting, we identified a conversion of the form *y = mx + b*. Because *b* was at most 11% of m, we approximated the calibration factor as *m*, which was then used to convert arbitrary fluorescence values to µM FITC (**Supplementary Figure 1**).

To characterize coacervate mixture, IVT, or ROSALIND fluorescence, 19 µLs of each sample was loaded onto a 384-well, optically clear, flat-bottomed plate covered with a plate seal and placed into a plate reader (Synergy H1, BioTek Gen5 v.3.10). Kinetic analysis of 3WJdB fluorescence was performed by reading the plate at 3-min intervals with excitation and emission wavelengths of 472 and 507 nm, respectively, for 2-4 hours. Freshly assembled (i.e., non-lyophilized) reactions were shaken in the plate reader for 10 s every 3 min during the 2-4 hour read period. Rehydrated lyophilized reactions were not shaken inside the plate reader to better approximate field conditions. Arbitrary fluorescence values were then converted to µM FITC by dividing by the appropriate calibration factor. Time to visible signal was determined by identifying the minutes elapsed until a fluorescence value of at least 1 µM FITC was reached.

### Hill equation fits

Where indicated, data were fit to the Hill equation with the functional form

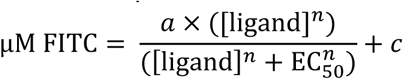

where [ligand] denotes ligand concentration, *c* represents the response with no ligand, *a* represents the maximum response, and *n* describes the cooperativity. The fitting was performed by Datagraph 5.2 on the average of the replicates, with initial parameterization at *a* = 1, EC_50_ = 5, *c* = 1, and *n* = 1. For the dose-response curves, the EC_50_ values were obtained from the fits generated by Datagraph 5.2. The limits of detection (LOD) and limits of quantification (LOQ) values were determined by numerically inverting the Hill Equation and computing the ligand concentration corresponding to 3 and 10 standard deviations above the blank mean, respectively. The visible concentration (VC), defined as the analyte concentration necessary to induce visible signal in 2 hours, was determined by numerically inverting the Hill Equation and computing the ligand concentration corresponding to 1 µM FITC signal.

## CODE AVAILABILITY

The Jupyter Notebook files with the Python codes used to compute the LOD, LOQ, and VC are provided in **Supplementary Data 4**.

## DATA AVAILABILITY

All data presented in this paper are available as **Source Data** and as **Supplementary Information**. A table of DNA sequences, along with Addgene accession numbers, is provided in **Supplementary Data 1**. A list of supplementary data files is included in the **Supplementary Information**. Additional materials, such as raw microscopy images, are available on the Zenodo open repository (DOI: 10.5281/zenodo.20648254).

## Supporting information

Jupyter Notebook File

## ACKNOWLEDGEMENTS

We thank K. Bhagat (Northwestern University) and A. Moreno (Northwestern University) for managing the experimental reagents and equipment used in this study; Y. Li (Northwestern University) for helpful discussions on ROSALIND and TMSD circuits; T.J. Lucci (Northwestern University) for providing purified smtB for preliminary experiments and helpful discussions on lyophilization; J. Fu (Northwestern University) for helpful discussions on RNA purification and quantification; A. Holkar (University of California, Los Angeles) for helpful discussions on coacervate design and selection; V. Trada (University of California, Los Angeles) for helpful discussions on coacervate properties; C. Fedosejevs (Northwestern University) for helpful discussions on microscopy; K. Shrinivas (Northwestern University) for helpful discussions on biological condensation; and N.P. Kamat (Northwestern University) and R. Schulman (Johns Hopkins University) for helpful discussions on compartmentalization of cell-free reactions. S.F. was supported by the National Science Foundation Research Traineeship (2021900) through Northwestern University’s Synthesizing Biology Across Scales Training Program. A.G. was supported by the UC Santa Barbara Chancellorʹs Postdoctoral Fellowship Program. This work was also supported by the Army Research Office (W911NF-22-2-0246) and by Army Contracting Command (W52P1J-21-9-3023), both to J.B.L., and by the National Science Foundation (DMR 2048285), to S.S.. This work made use of the Northwestern University SynBio Foundry Core Facility (RRID:SCR_026869). This work was also supported by the BioPACIFIC Materials Innovation Platform of the National Science Foundation (DMR-2445868). The views, opinions, and/or findings expressed are those of the authors and should not be interpreted as representing the official views or policies of the Department of Defense, the National Science Foundation, or the U.S. Government.

## AUTHORS AND AFFILIATIONS

Department of Biomedical Engineering, Northwestern University, Evanston, IL, USA Siyuan Feng

Center for Synthetic Biology, Northwestern University, Evanston, IL, USA Siyuan Feng, Rebecca A. Rasmussen, Lauren Clark & Julius B. Lucks

Department of Chemical and Biological Engineering, Northwestern University, Evanston, IL, USA

Rebecca A. Rasmussen & Julius B. Lucks

California Nanosystems Institute, University of California, Santa Barbara, Santa Barbara, CA, USA

Antonio Garcia IV

Department of Chemical and Biomolecular Engineering, University of California, Los Angeles, Los Angeles, CA, USA.

Antonio Garcia IV & Samanvaya Srivastava

BioPACIFIC Materials Innovation Platform, University of California, Los Angeles, Los Angeles, CA, USA.

Samanvaya Srivastava

California NanoSystems Institute, University of California, Los Angeles, Los Angeles, CA, USA. Samanvaya Srivastava

Institute for Carbon Management, University of California, Los Angeles, Los Angeles, CA, USA. Samanvaya Srivastava

Interdisciplinary Biological Sciences Graduate Program, Northwestern University, Evanston, IL, USA

Julius B. Lucks

Center for Water Research, Northwestern University, Evanston, IL, USA Julius B. Lucks

Center for Engineering Sustainability and Resilience, Northwestern University, Evanston, IL, USA

Julius B. Lucks

## Contributions

S.F., S.S., and J.B.L. designed the study. S.F., S.S., and J.B.L. analyzed the data. S.F., R.A.R., A.G., and L.C. conducted the research. S.F., A.G., S.S., and J.B.L. developed the methodology. S.F. and J.B.L. undertook data visualization. S.F. and J.B.L. curated the data. S. S. and J.B.L. acquired funding for the study. S.F. and A.G. validated the results. S.F. and J.B.L. managed and coordinated the study. S.F., S.S., and J.B.L. supervised the research and wrote the manuscript. All authors edited the manuscript.

## Corresponding author

Correspondence to Samanvaya Srivastava and Julius B. Lucks.

## ETHICS DECLARATIONS

### Competing interests

S.F., S.S., and J.B.L. are inventors on a US patent application (63/797,755) relating to the use of coacervates to enhance cell-free biosensing. S.S. is an inventor on a US application (18/559,983) on coacervate bioreactors. J.B.L. is an inventor on US patents (US12098433, US12203144, US12325884) and is an inventor on US applications (17/606,441; 19/229,877; 18/548,175; 18/311,030; 19/161,065; 63/625,752; 63/637,174; 63/805,130; 63/866,893; 63/755,965; 63/705,786) on cell-free biosensing technologies. R.A.R., A.G., and L.C. declare no competing interests.

## Notes

https://zenodo.org/records/20648254

