## Supplementary Information for "Transcription-induced coacervation accelerates and sensitizes cell-free biosensing"

### SUPPLEMENTARY METHODS

**Charge concentration calculations.** Charge concentration is conventionally used in coacervates research to describe polyelectrolyte concentration, as these polymers can differ substantially in length and thus in the number of charges carried per molecule and in their contribution to charge balance. 200 mM charge concentration PAA (5100 g/mol) describes the amount of PAA added that would contribute 200 mM of charge, assuming (1) a polydispersity index of 1 and (2) each repeating unit is fully ionized and contributes charge -1. Under these idealized conditions, each PAA molecule contains 54.26 of charge, determined by dividing the molecular weight of the polymer (5100 g/mol) by the molecular weight of the repeating unit (94 g/mol). Therefore, 200 mM charge concentration PAA corresponds to 3.7 mM PAA or 0.38 mg of PAA per 20 µL reaction.

**T7 RNAP expression, purification, and labeling.** T7 RNAP was expressed and purified as described previously [1], with minor modifications, to generate RNAP to label for microscopy. Purification buffers containing Tris base were replaced with MOPS and buffered to pH 7.5 to be compatible with NHS ester labeling. Briefly, BL21(DE3) cells were transformed with plasmid pAR1219 (Sigma-Aldrich). 4 Ls of cell culture were grown in 2X YTPG medium at 37 °C, induced with 1 mM IPTG at an OD600 of 0.6, and grown for four additional hours (approx. OD600 ≈ 10) at 37 °C. The cultures were pelleted by centrifugation, flash frozen, and stored at −70 °C. On the day of purification, pellets were thawed, resuspended in a lysis buffer (50 mM HEPES, 20 mM NaCl, 2 mM EDTA, 1 mM DTT) with HALT protease inhibitor, and homogenized using an Avestin C5 homogenizer. Ammonium sulfate followed by polymin P was added to the lysed cells to precipitate nucleic acids, and the insoluble materials were removed by centrifugation. Proteins were then precipitated by adding 4.1 M ammonium sulfate pH 7 with KOH, collected through centrifugation, and the pellet was resuspended in glycerol-free Buffer C (20 mM potassium phosphate pH 7.7, 1 mM EDTA, 1 mM DTT) with 100 mM NaCl. Glycerol-free buffers were used to enhance compatibility with downstream NHS ester labeling. After overnight dialysis, the sample was filtered through a 0.2 µm filter and purified through cation exchange chromatography using a SP-Sepharose column (Cytiva HiPrep SP FF 16/10) on an AKTA Avant 25. The column was washed with 4 column volumes of glycerol-free Buffer C with 50 mM NaCl, and T7 RNAP was eluted with glycerol-free Buffer C with 200 mM NaCl. Fractions were analyzed by SDS–PAGE, and those containing T7 RNA polymerase were pooled and dialyzed twice against glycerol-free Buffer C with 10 mM NaCl for 2 hours each at 4 °C. The sample was then dialyzed overnight against MOPS-based S30 buffer (10 mM MOPS, pH 7.5, 14 mM magnesium acetate, and 60 mM potassium acetate) with 2 mM DTT. Samples were centrifuged to remove precipitates, and the concentration of the protein in the supernatant was determined by protein Qubit.

**aTF purification.** aTFs were expressed and purified as described previously [2], with minor modifications. The protein expression plasmids (**Supplementary Data 1**) were transformed into BL21(DE3) competent cells. 1-4 Ls of cell culture were grown by shaking at 250 RPM at 37 °C until OD600 ≈ 0.5, at which point 0.5 mM IPTG was added. Cultures remained at 37 °C with 250 RPM shaking for an additional 4 hr. Cultures were pelleted by centrifugation at 4000 xg at 4 °C for at least 20 minutes. Pellets were flash-frozen in liquid nitrogen and stored at -70 °C until further processing.

The frozen pellets were thawed slowly on ice and then each gram of pellet was resuspended in 5-10 mL of lysis buffer (10 mM Tris-HCl pH 8, 500 mM NaCl, 1 mM TCEP-HCl, 1 protease inhibitor tablet (cOmplete EDTA-free Protease Inhibitor Cocktail, Roche) per 50 mL). The resuspended pellets were lysed using a Q125 sonicator for 5 cycles of 15 seconds on, 45 seconds off at 50% amplitude. The lysates were centrifuged at 13000 xg at 4 °C for 30 min to remove insoluble debris. The soluble lysate was then filtered with 0.22 μm filters.

The proteins TetR, SmtB and CsoR were purified with affinity chromatography in a gravity column. The column was filled with 4 mL of Ni-NTA agarose (Qiagen) and equilibrated with 10 mL of bind buffer (10 mM Tris-HCl pH 8, 500 mM NaCl, 1 mM TCEP-HCl) with 20 mM imidazole. The filtered supernatant was passed through the column, followed by 2 additional wash steps with 10 mL bind buffer with 20 mM imidazole. Except for CsoR purification, the column was washed and eluted with 5 mL fractions of increasing imidazole concentration in bind buffer: 20-50 mM in increments of 5 mM, 50-100 mM in increments of 10 mM, 150 mM, and 300 mM. CsoR was eluted through 10 mL fractions of bind buffer with 25, 30, 35, 40, 45, and 300 mM imidazole. The fractions were assessed using SDS-PAGE. CsoR was dialyzed overnight into TEV cleavage buffer (50 mM Tris pH 8, 250 mM NaCl, 5 mM DTT), and the 6x His-tag from CsoR was removed using TEV protease (New England Biolabs) by incubating at 30 °C for 1 hr and overnight at 4 °C. The cleavage reactions were separated using the gravity nickel columns, eluting with bind buffer plus 50, 100, 150, 200, and 300 mM imidazole. These fractions were assessed using SDS-PAGE.

The proteins MobR and TtgR were purified using fast protein liquid chromatography using nickel affinity chromatography followed by gel filtration on an AKTA Avant 25. The HisTrap FF, 5 mL column equilibrated with bind buffer (10 mM Tris-HCl pH 8, 500 mM NaCl, 1 mM TCEP-HCl). The filtered protein expression lysate was applied at 5 mL/min. The column was then washed with bind buffer with 20 mM imidazole at 5 mL/min, collecting 2 mL fractions. The imidazole concentration was then gradually increased to 300 mM, while maintaining the 5 mL/min flow rate and 2 mL fraction collection. Selected based on 280 nm absorbance peaks, fractions were analyzed by SDS-PAGE. The fractions of interest were buffer exchanged in a 10 MWCO or dialyzed in a 6-8 MWCO dialysis bag into bind buffer.

The MobR and TtgR protein samples were then spin concentrated in a 10 MWCO Amicon centrifugal filter to load into a 2 mL loop for gel filtration. A HiLoad 16/600 Superdex 200 pg column was equilibrated with bind buffer. The sample was loaded from the loop and eluted with bind buffer with 1 mL/min flow, collecting 2 mL fractions. Fractions with an increased absorbance at 280 nm were analyzed by SDS-PAGE.

Fractions of pure protein from the gravity affinity purification or FPLC gel filtration were combined and buffer exchanged 3 times in a 10 MWCO Amicon with 10 mL 2x storage buffer (50 mM Tris-HCl pH 8, 200 mM NaCl, 4 mM TCEP), modified to 100 mM Tris for CsoR, SmtB, and TetR. The proteins were further concentrated to ~100 to 200 μL. Equal volume of 100% glycerol was added and mixed to create a 50% glycerol stock for each protein. The concentrations of the protein samples were measured using protein Qubit.

**Supplementary Table 1 | Polymer identity and concentrations in tested candidate coacervate systems.**

| Coacervate System | Polycation Molecular Weight (g/mol) | [Polycation] (mM) | Polycation charge concentration (mM) | Polyanion Molecular Weight (g/mol) | [Polyanion] (mM) | Polyanion charge concentration (mM) |
|---|---|---|---|---|---|---|
| **PDADMAC-PAA** | poly(diallyldimethyl ammonium chloride) 8500 | 0.38 | 20 | poly(acrylic acid sodium salt) 5100 | 0.37 | 20 |
| **spermidine -PAA** | spermidine trihydrochloride 254.63 | 2[a] | 6[a] | poly(acrylic acid sodium salt) 5100 | 0.37 | 20 |
| **spermine-PAA** | spermine 202.34 | 4 | 16 | poly(acrylic acid sodium salt) 5100 | 0.37 | 20 |

[a] This concentration is in addition to the 2 mM spermidine already present in transcription buffer, within which all three candidate systems were assembled and tested.

**Supplementary Table 2 | Transcription buffer components.**

| Component | Concentration (mM) | Purchased From |
|---|---|---|
| Tris-HCl, pH 8 | 40 | Invitrogen (AM9856) |
| MgCl2 | 8 | Sigma-Aldrich (M8266) |
| Dithiothreitol | 10 | GoldBio (DTT100) |
| NaCl | 20 | Sigma-Aldrich (S3014) |
| Spermidine | 2 | ThermoScientific (215100050) |

**Supplementary Table 3 | ROSALIND component and ligand concentrations.** All aTF concentrations are listed as dimer concentrations, except for CsoR, which is a tetramer.

| Ligand-aTF Pair | [DNA] (nM) | [aTF] (μM) | [Ligand] (μM) | Ligand Solvent | [Ligand Stock] (μM) | Ligand Purchased From |
|---|---|---|---|---|---|---|
| Anhydrotetracycline HCl - TetR | 25 | 0.15 | 0.18 - 1.93 | Ethanol | 233.3 | Sigma-Aldrich (37919) |
| Tetracycline HCl - TetR | 25 | 0.15 | 0.2 - 40 | Water | 138.6 | GoldBio (T-101-25) |
| ZnSO4 - SmtB | 25 | 6 | 0.5 - 75 | Water | $2 \times 10^6$ | Sigma-Aldrich (83265) |
| CuSO4 - CsoR | 100 | 4.5 | 1 - 41 | Water | $1 \times 10^5$ | Sigma-Aldrich (1.02784) |
| 3-OH Benzoic Acid - MobR | 25 | 32 | 1.5 - 500 | 1 M Tris pH 8 | $1 \times 10^5$ | Sigma-Aldrich (H20008) |
| Naringenin - TtgR | 25 | 10 | 1 - 250 | Ethanol | $1 \times 10^4$ | Sigma-Aldrich (N5893) |
| ZnSO4 - SmtB (reduced DNA) | 10 | 2.5 | 0.5 - 75 | Water | $2 \times 10^6$ | Sigma-Aldrich (83265) |
| ZnSO4 - SmtB (lyophilized, lab water) | 25 | 6 | 0.5 - 75 | Water | $2 \times 10^6$ | Sigma-Aldrich (83265) |
| ZnSO4 - SmtB (lyophilized, lake water) | 25 | 18 | 0.83 - 75 | Water | $2 \times 10^6$ | Sigma-Aldrich (83265) |

**Supplementary Table 4 | Sample preparation for confocal imaging.**

| Imaging sample | Figure | AlexaFluor546-DNA (nM) | Dylight405-RNAP (µg) | PAA (charge mM) | spermine (mM) |
|---|---|---|---|---|---|
| **One-color (yellow)** | SI Figure 8 | 25 | 0 | 20 | 4 |
| **One-color (blue)** | SI Figure 9 | 0 | 8.6 | 20 | 4 |
| **Two-color** | Figure 3C | 12.5 | 4.3 | 20 | 4 |

**Supplementary Table 5 | Purification methods for aTFs.**

| aTF | Type of Purification | Tag Location | TEV Cleavage | Columns Used |
|---|---|---|---|---|
| **TetR** | His-tag affinity | C-terminal 6x His-tag | No | Gravity flow column with Qiagen Ni-NTA Agarose |
| **SmtB** | His-tag affinity | C-terminal TEV cleavage site followed by 6x His-tag | No | Gravity flow column with Qiagen Ni-NTA Agarose |
| **CsoR** | His-tag affinity | N-terminal 6x His-tag followed by TEV cleavage site | Yes | Gravity flow column with Qiagen Ni-NTA Agarose |
| **MobR** | His-tag affinity followed by size exclusion chromatography | C-terminal TEV cleavage site followed by 6x His-tag | No | Cytiva HisTrap FF 5mL nickel column for affinity, Cytiva Superdex HiLoad 16/600 200 pg column for size exclusion |
| **TtgR** | His-tag affinity followed by size exclusion chromatography | C-terminal 6x His-tag | No | Cytiva HisTrap FF 5mL nickel column for affinity, Cytiva Superdex HiLoad 16/600 200 pg column for size exclusion |

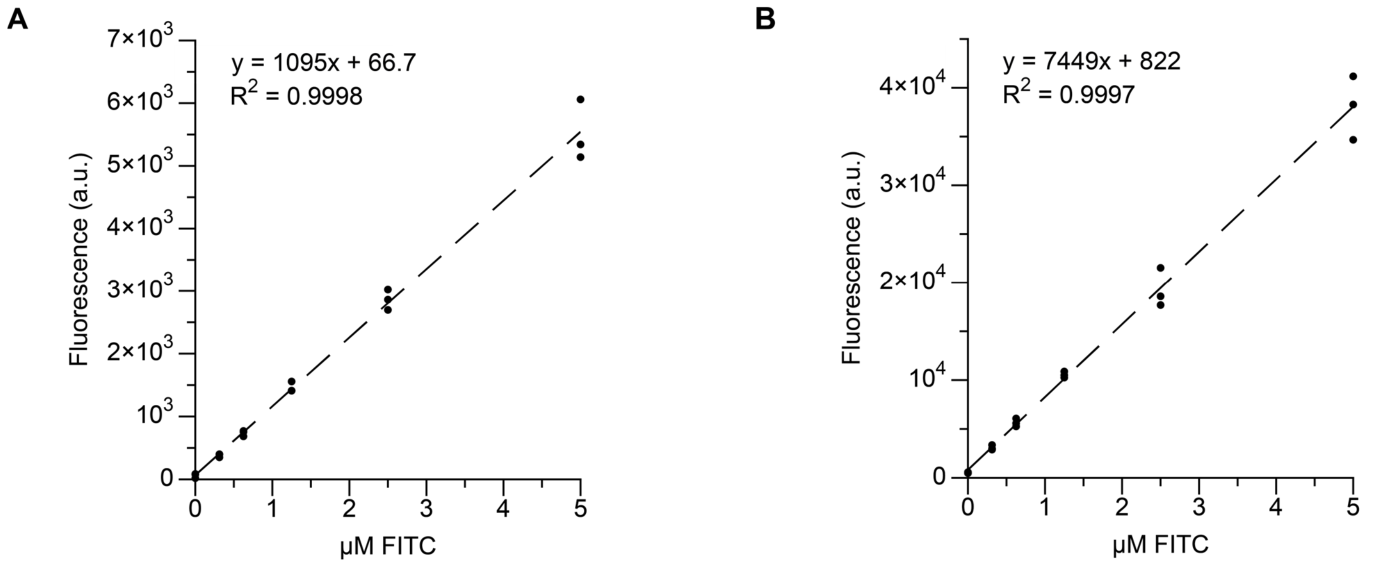

**Supplementary Fig. 1 | Fluorescence standardization to micromolar fluorescein (µM FITC).** Arbitrary units of fluorescence were standardized to µM concentrations of fluorescein using a NIST traceable standard (see **Methods**). In the representative examples shown here, a dilution series of FITC standard was prepared in buffer (100 mM sodium borate, pH 9.5), and measured on a plate reader using the same settings for measuring (**A**) 3WJdB (472 nm excitation, 507 nm emission) signal or (**B**) 6-FAM (495 nm excitation, 520 nm emission) signal. The resulting data were then used to standardize fluorescence measured from ROSALIND reactions. The standard curve was generated for each plate reader and each measurement setting. Data are shown for n=3 replicates of each sample.

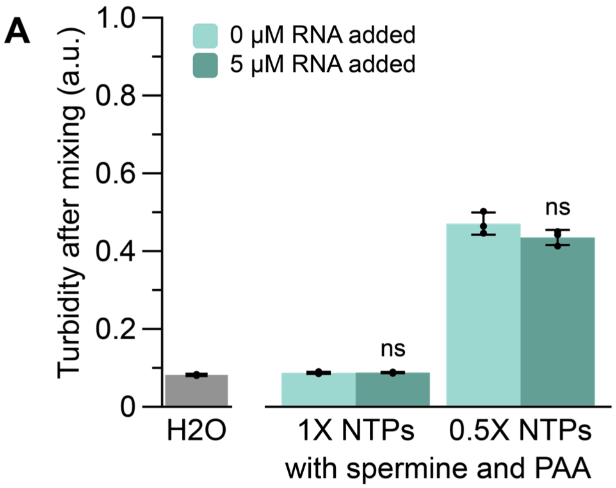

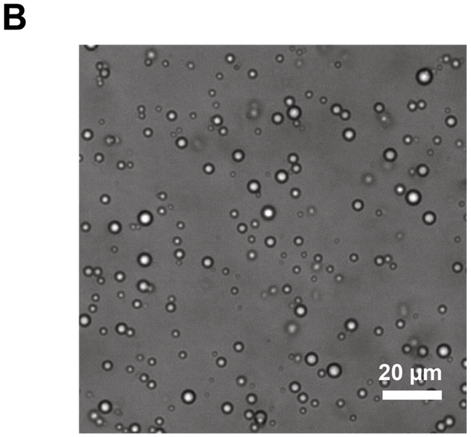

**Supplementary Fig. 2 | Addition of RNA without NTP consumption does not induce or alter coacervation.** 5 µM purified RNA was added to IVT reactions with spermine and PAA, without T7 RNAP to preclude NTP consumption. **(A)** Addition of RNA does not induce or increase turbidity in reactions assembled with full or partially depleted (0.5X) NTPs. Reactions were assembled on ice with turbidity measurements taken in a plate reader set to room temperature. Bar heights represent the average of the n=3 independent replicates, and the error bars indicate the average value of the replicates +/- the standard deviation. A two-tailed, heteroscedastic Student's *t*-test was used to compare samples with and without RNA at each NTP concentration; ns denotes that the P value is greater than 0.05. **(B)** Representative image taken of an IVT reaction without T7 RNAP, with 0.5X NTPs and 5 µM RNA immediately after mixing. The coacervates that form resemble those that are formed under these conditions without the addition of RNA (**Figure 2E**).

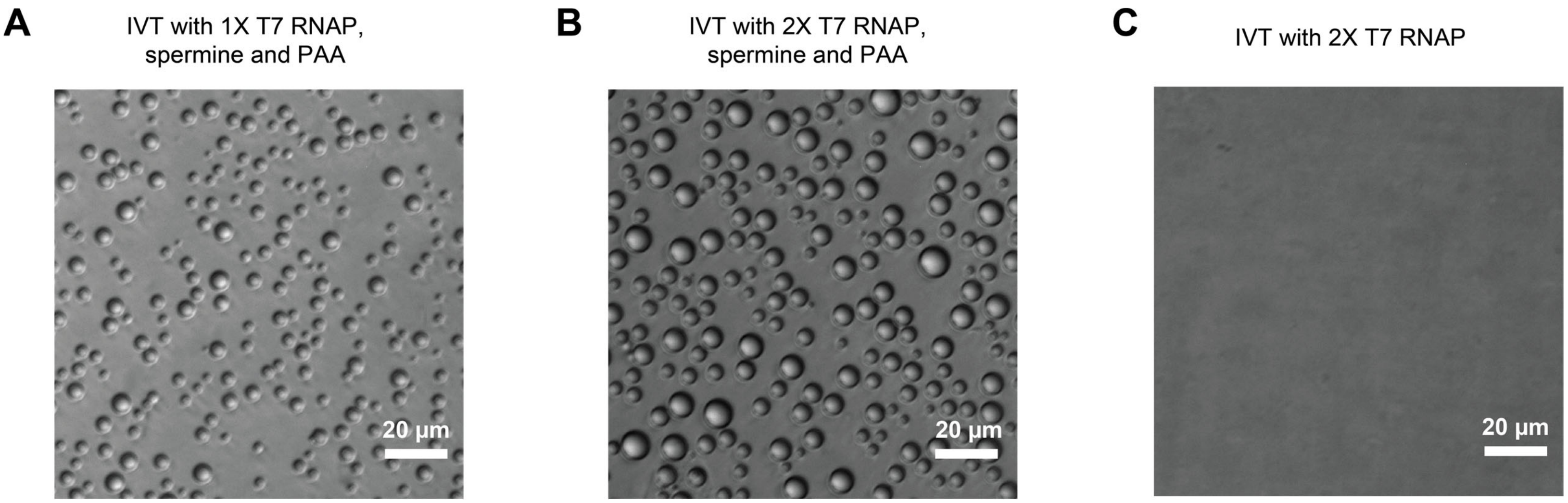

**Supplementary Fig. 3 | Enhancing IVT by increasing RNAP concentration leads to greater coacervate droplet number and size.** IVT reactions containing spermine and PAA were assembled with either **(A)** 10 U $\mu L^{-1}$ (1X) or **(B)** 20 U $\mu L^{-1}$ (2X) T7 RNAP and imaged with DIC microscopy after 4 hours of incubation at 37 ºC. IVT reactions with more T7 RNAP showed greater droplet number and size. **(C)** A control IVT reaction assembled with 20 U $\mu L^{-1}$ (2X) T7 RNAP without spermine or PAA does not show droplets under DIC microscopy after 4 hours of incubation at 37 ºC.

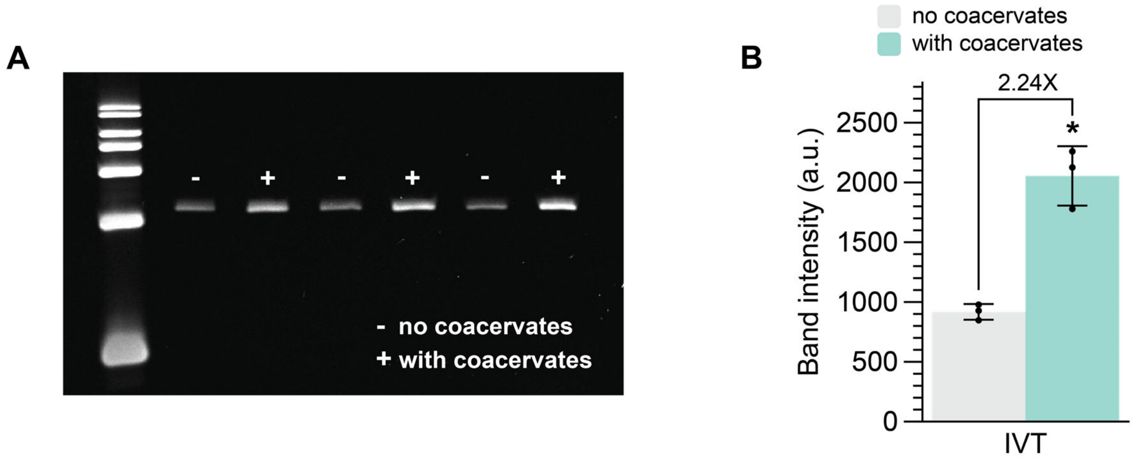

**Supplementary Fig. 4 | Quantification of RNA urea-PAGE gel shows that coacervates increase RNA production by 2X.** IVTs were incubated for 2 hours at room temperature, and the transcribed RNA was isolated after DNAse digest with Trizol extraction. Samples were run on a 7% urea-PAGE gel, shown in **(A)**, and the band intensity was quantified using ImageJ/FIJI, shown in (**B**). A two-tailed, heteroscedastic Student's *t*-test was used to compare samples with and without coacervates; * denotes that the P value is between 0.01 and 0.05. Bar heights represent the average of the n=3 independent replicates, and the error bars indicate the average value of the replicates +/- the standard deviation. The uncropped, unprocessed gel image is available in the **Source Data**.

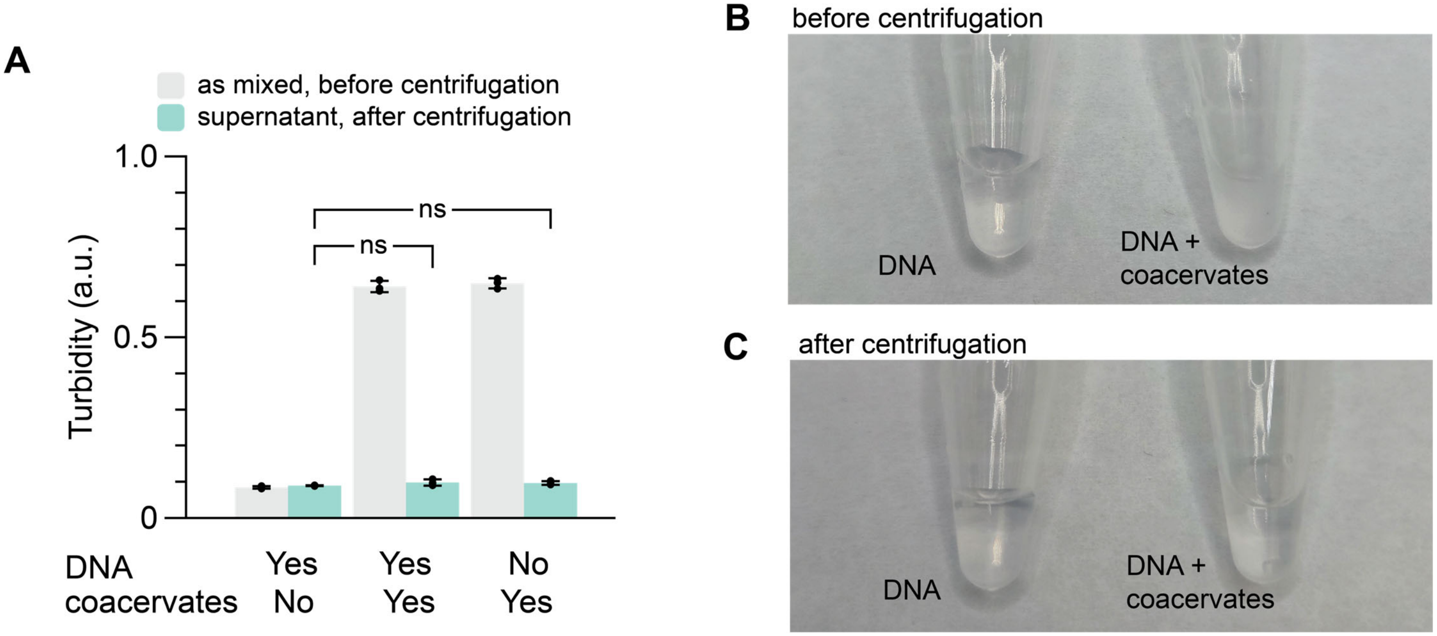

**Supplementary Fig. 5 | Centrifugation separates the dense coacervate phase from the dilute supernatant.** Samples containing DNA, DNA with spermine-PAA coacervates, or spermine-PAA coacervates alone were centrifuged for 5 mins at 5000 RCF. **(A)** Turbidity measurements before and after centrifugation indicate that centrifugation successfully pellets the coacervate phase and removes droplets. After centrifugation, the turbidity of samples with coacervates is statistically indistinguishable from that of samples without coacervates. **(B-C)** Representative images of the samples are shown before and after centrifugation. Samples with DNA without coacervates are optically clear prior to centrifugation and their turbidity does not change after centrifugation. Samples with DNA and coacervates are opaque prior to centrifugation and become clear after centrifugation. Bar heights in **(A)** represent the average of the n=3 independent replicates, and the error bars indicate the average value of the replicates +/- the standard deviation. Significance was determined using a two-tailed, heteroscedastic Student's t-test against the turbidity of the DNA-only sample after centrifugation; ns denotes that the P value is greater than 0.05. The uncropped, unprocessed images in **(B-C)** are available in the **Source Data**.

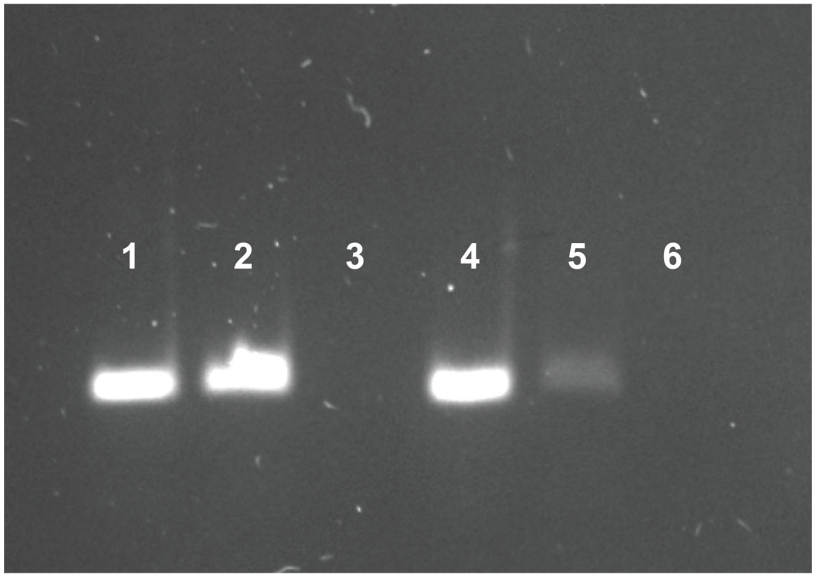

**Supplementary Fig. 6 | Agarose gel electrophoresis of centrifuged samples shows that DNA is sequestered by the dense coacervate phase.** Samples containing DNA, DNA with spermine-PAA coacervates, or spermine-PAA coacervates alone, were centrifuged to pellet the dense coacervate phase. Electrophoresis lanes were loaded with as-mixed samples collected prior to centrifugation from mixtures containing DNA (lane 1), DNA with spermine-PAA coacervates (lane 2), or spermine-PAA coacervates alone (lane 3), or with samples collected from the supernatant from these same mixtures after centrifugation (lanes 4-6, in the same order). The difference between lane 2 and lane 5 suggests that the DNA is sequestered by the dense coacervate phase. The uncropped, unprocessed image corresponding to these samples and two additional replicates are available in the **Source Data** for **Figure 3**.

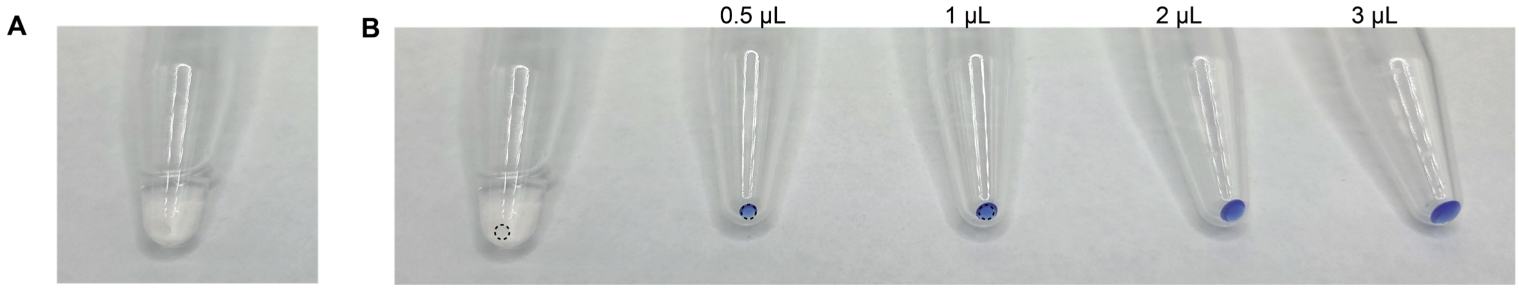

**Supplementary Fig. 7 | The coacervate phase after centrifugation is approximately 1 μL. (A)** After centrifugation, the coacervate phase forms a small, clear pellet at the bottom of the tube. **(B)** Following a published method [3], we compared the coacervate pellet with a set of known volumes of organic dye, which suggests that the dense pelleted coacervate phase is approximately 1 μL. Because the dense phase can be compressed during centrifugation due to water loss, this packed volume may underestimate the true volume occupied by the coacervates when dispersed. In other systems, dense coacervate phases contain 50–75% water [4, 5]; assuming maximal water loss during centrifugation, the uncompressed coacervate phase volume is likely at or below 5 μL, or 5% of the system volume (100 μL). Results are shown for representative images. The image in **(A)** is a cropped duplicate of **(B)**. The black guidelines in **(B)** are of the same size superimposed over the coacervate pellet and the 0.5 μL and 1 μL volume standards. The organic dye is 0.024% bromophenol blue dissolved in formamide. The uncropped, unprocessed images are available in the **Source Data**.

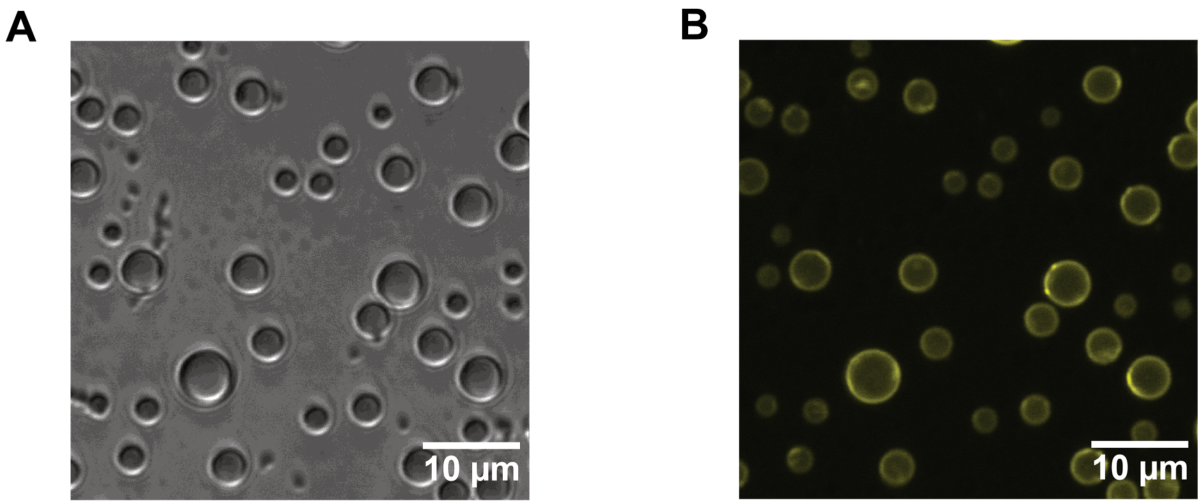

**Supplementary Fig. 8 | DNA localizes to the coacervate dense phase, especially to the liquid interface. (A)** Brightfield and **(B)** confocal images of spermine-PAA coacervates with Alexa Fluor 546-labeled DNA. These samples were prepared as described in **Supplementary Table 4**.

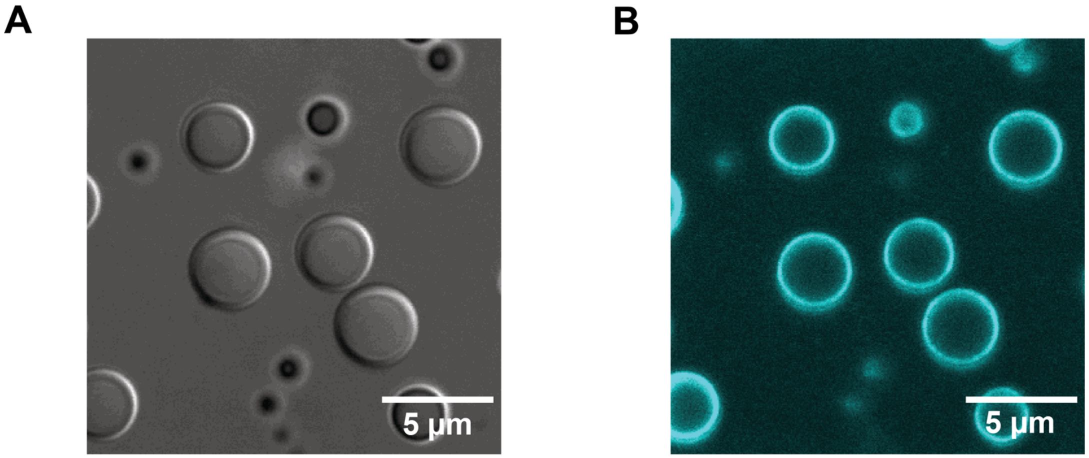

**Supplementary Fig. 9 | T7 RNAP localizes to the coacervate dense phase, especially to the liquid interface. (A)** Brightfield and **(B)** confocal images of spermine-PAA coacervates with DyLight 405-labeled T7 RNAP. These samples were prepared as described in **Supplementary Table 4.**

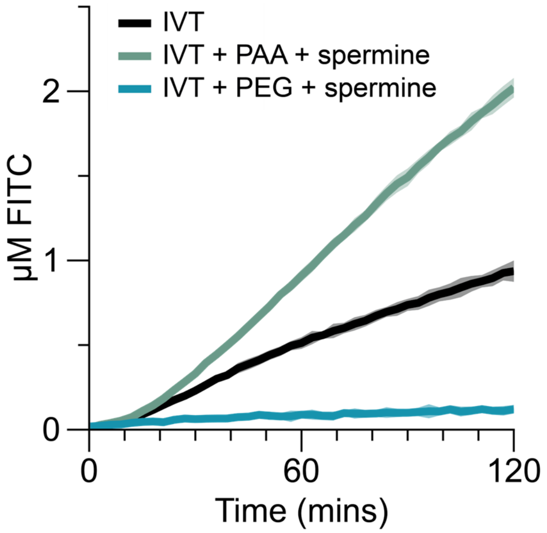

**Supplementary Fig. 10 | Replacing PAA with an equiweight concentration of PEG results in transcriptional inhibition.** The addition of PEG and spermine to IVT reactions transcribing 3WJdB at room temperature resulted in near ablation of reporter signal. PEG was added in equiweight concentration to the amount of PAA in spermine-PAA coacervates (**Supplementary Table 1**). The control line shown (in black) is the same as the line in **Figure 3D**, and is added to facilitate comparison with a control IVT reaction. Shading indicates the average value of n=3 independent biological replicates +/- standard deviation. Raw fluorescence values were standardized to µM FITC.

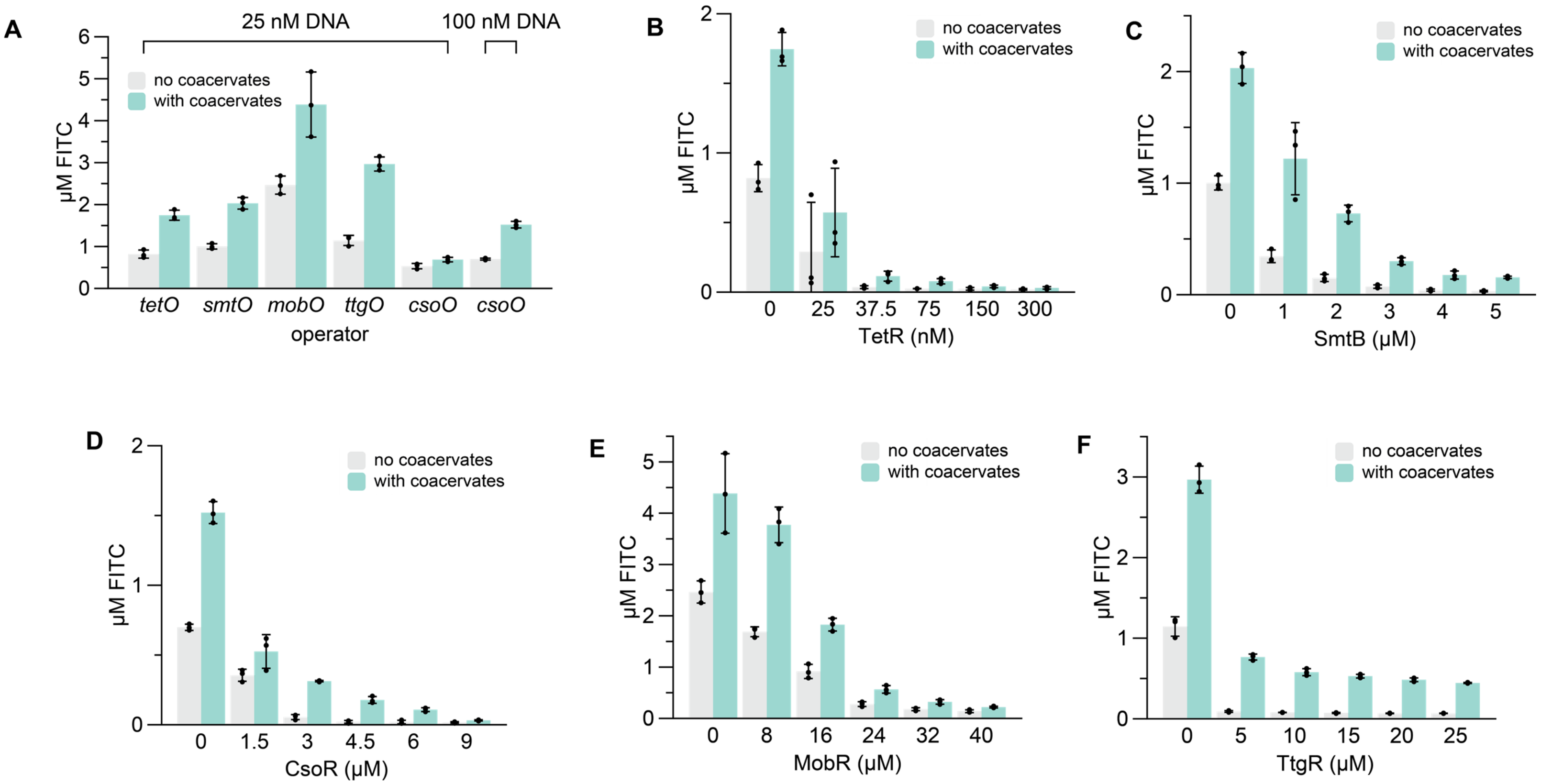

**Supplementary Fig. 11 | Coacervates are compatible with modular repression by aTFs. (A)** Spermine-PAA coacervates enhance the unregulated transcription of DNA templates containing diverse aTF operators. Transcription of the *csoO* template results in weak reporter output (0.53 µM FITC after 2 hours), and the smallest coacervate-mediated enhancement (1.3-fold, to 0.69 µM FITC). We hypothesized that weaker transcription from this template hampers coacervate formation and thus limits the enhancement effect. To test this hypothesis, we strengthened transcription by increasing the DNA concentration from 25 nM to 100 nM, and observed an increase in the coacervate-mediated improvement from 1.3-fold to 2.1-fold in coacervate-mediated fold improvement in unregulated IVT reactions containing *csoO* template. We chose to use 100 nM DNA in our subsequent experiments involving the *csoO* operator. **(B)** TetR can be used with coacervates to repress transcription from 25 nM DNA template containing the *tetO* operator. **(C)** SmtB can be used with coacervates to repress transcription from 25 nM DNA template containing the *smtO* operator. **(D)** CsoR can be used with coacervates to repress transcription from 100 nM DNA template containing the *csoO* operator. **(E)** MobR can be used with coacervates to repress transcription from 25 nM DNA template containing the *mobO* operator. **(F)** TtgR can be used with coacervates to repress transcription from 25 nM DNA template containing the *ttgO* operator. These aTFs span four unique families (TetR, ArsR/SmtB, CsoR/RcnR, and MarR). Data in **(A)** is repeated as the leftmost bar in panels **(B-F)** to facilitate comparison with the unrepressed transcription from the corresponding DNA template. All protein concentrations in **(B-F)** are dimer concentrations, except for CsoR, which is a tetramer. All reactions were run at room temperature. Bar heights represent the average of the n=3 independent replicates, and the error bars indicate the average value of the replicates +/- the standard deviation. Raw fluorescence values were standardized to µM FITC.

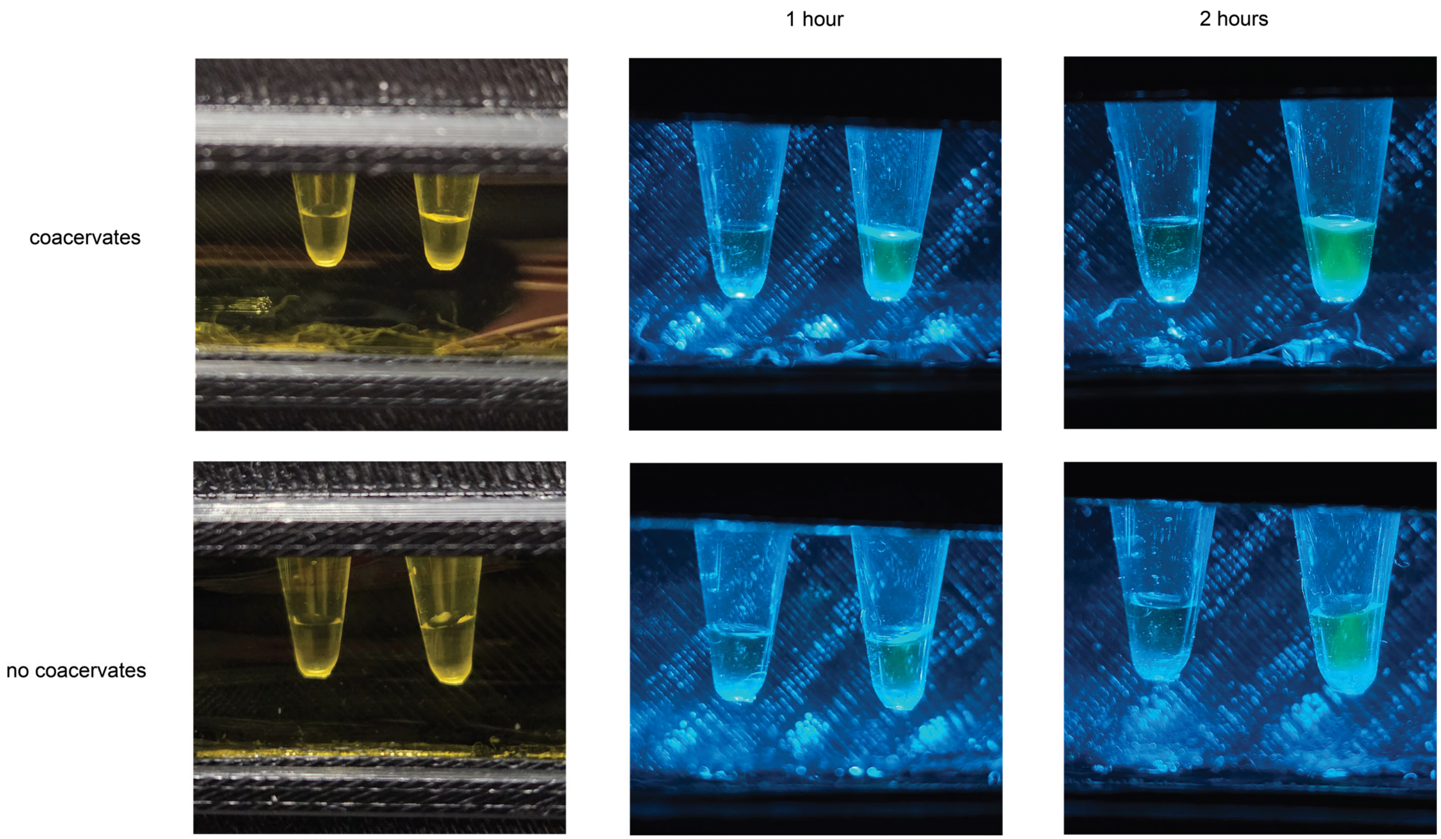

**Supplementary Fig. 12 | Coacervates accelerate the kinetics of anhydrotetracycline biosensing.** Representative images of anhydrotetracycline biosensing reactions are shown after 1 or 2 hours of incubation at room temperature. The left and right tubes in each photograph contain 0 μM or 1.9 μM anhydrotetracycline, respectively. PCR tubes were placed in a handheld 3D-printed blue-light illuminator built as described previously [2] and photographed with a Samsung S25 phone camera. The leftmost images show the biosensor reactions illuminated under room light for reference. Uncropped, unprocessed images are available in the **Source Data**.

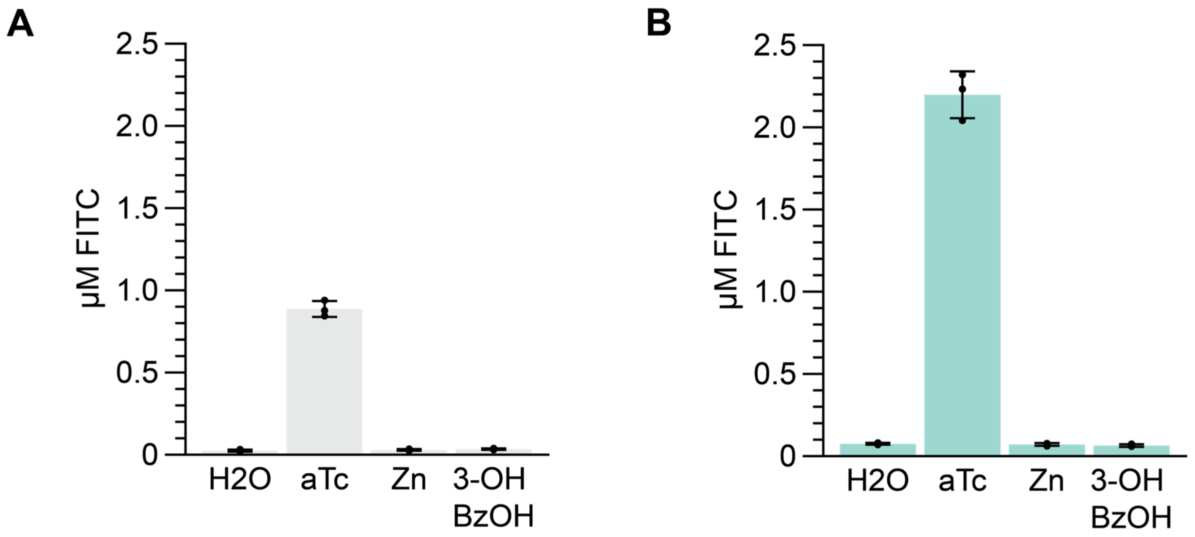

**Supplementary Fig. 13 | Coacervates preserve the specificity of TetR biosensing.** All reactions run at room temperature **(A)** without or **(B)** with spermine-PAA coacervates. H2O, aTc, Zn and 3-OH BzOH indicate that no ligand, 1.9 µM anhydrotetracycline, 75 µM zinc or 1 mM 3-hydroxybenzoic acid were added to the reaction. Bar heights represent the average of the n=3 independent replicates, and the error bars indicate the average value of the replicates +/- the standard deviation. Raw fluorescence values were standardized to µM FITC.

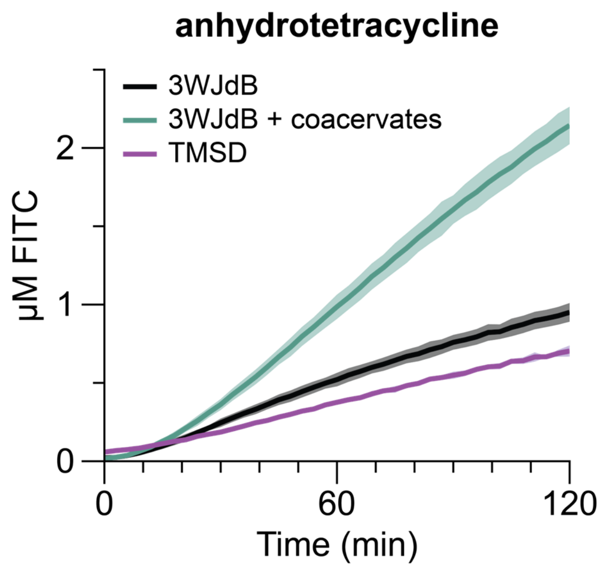

**Supplementary Fig. 14 | ROSALIND reactions with coacervates outperform reactions with TMSD at room temperature.** TMSD outputs are slower than 3WJdB outputs at room temperature, and reactions leveraging coacervates outperform those without coacervates and those leveraging TMSD at room temperature. All reactions contained 150 nM TetR dimer and 1.9 µM anhydrotetracycline and were run at room temperature. TMSD reactions contained 5 µM signal gate, the DNA duplex that undergoes strand displacement. Shading indicates the average value of n=3 independent biological replicates +/- standard deviation. Raw fluorescence values were standardized to µM FITC.

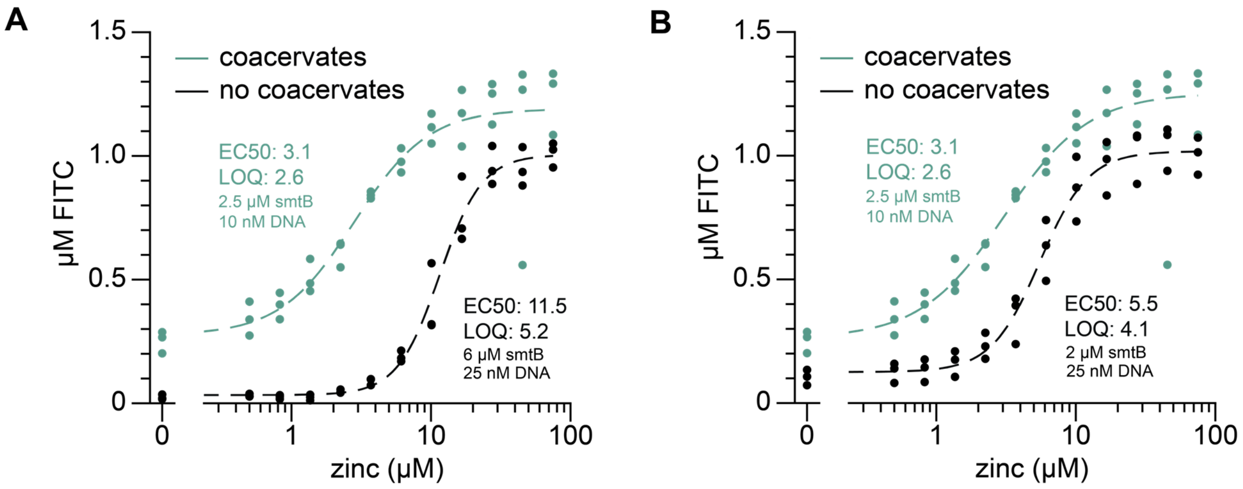

**Supplementary Fig. 15 | Coacervates can enable an additional sensitivity enhancement by reducing the DNA necessary to achieve signal.** ROSALIND reactions with 10 nM DNA and spermine-PAA coacervates achieve a similar ON signal at 2 hours to a ROSALIND reaction with 25 nM DNA without coacervates. Reactions with 10 nM DNA with coacervates can be repressed with 2.5 μM smtB. This enables a **(A)** 3.7-fold improvement and **(B)** 1.8-fold improvement in EC50 for zinc detection compared to control reactions with 25 nM DNA repressed with 6 μM or 2 μM smtB, respectively. The condition with coacervates is duplicated across **(A,B)** to facilitate visual comparison, and the control line in **(A)** is duplicated from **Figure 5C**. All reactions were run at room temperature. The smtB concentrations listed are dimer concentrations. Raw fluorescence values were standardized to μM FITC.

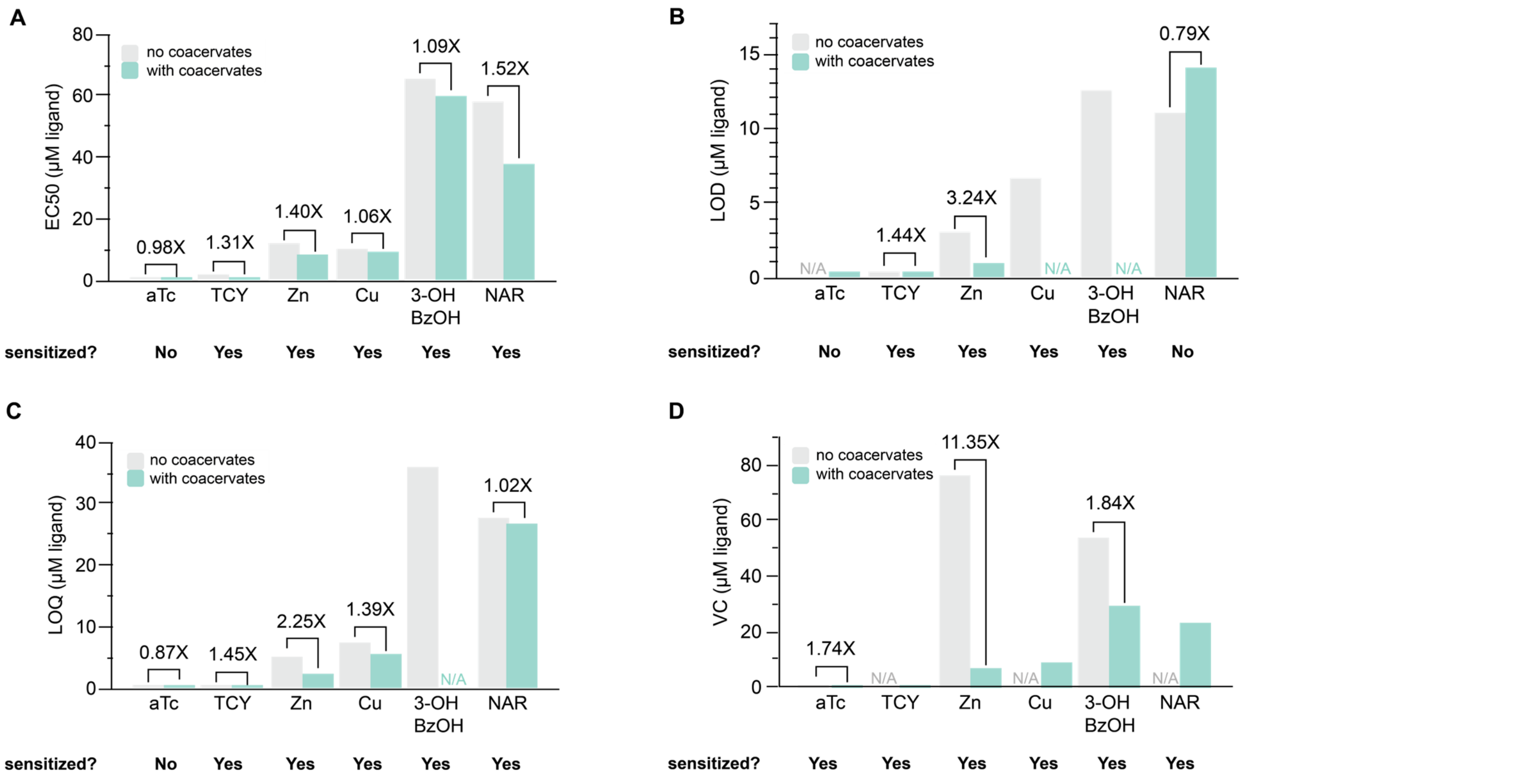

**Supplementary Fig. 16 | Comparison between EC50, limit of detection (LOD), limit of quantification (LOQ), and visible concentration (VC) sensitivity analyses.** Sensitivity metrics EC50 **(A)**, limits of detection **(B)**, limits of quantification **(C)**, and visible concentration **(D)** showed generally similar sensitization trends. The EC50 value describes the analyte concentration necessary to generate half-maximal signal. The LOD and LOQ values correspond to the analyte concentrations required to generate signals that are 3 or 10 standard deviations above the blank mean, respectively. The VC values denote the analyte concentration necessary to generate visible signal (1 µM FITC) in 2 hours. All sensitivity metrics were calculated from the dose-response curves shown in **Figure 5**. The EC50 values are the same as those noted in **Figure 5**. Fold changes are calculated by dividing the sensitivity metric without coacervates by that with coacervates. N/A describes conditions where the LOD, LOQ, or VC could not be calculated from the dose-response curve; additional information is provided in **Supplementary Data 3** and **Supplementary Data 4**.

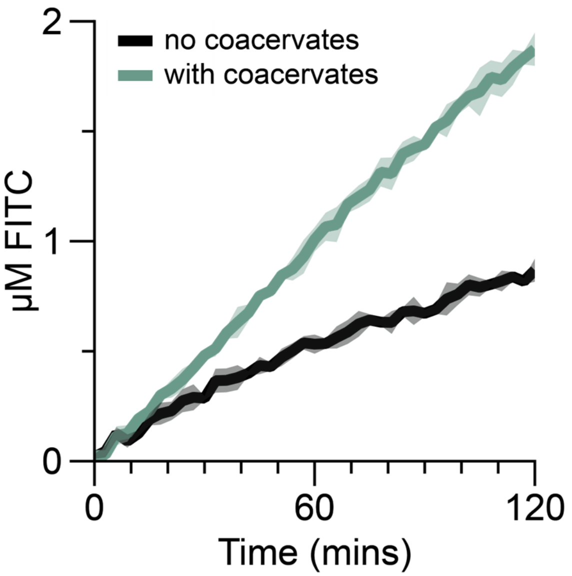

**Supplementary Fig. 17 | Rehydration of freeze-dried unregulated IVT reactions with coacervates demonstrate improved kinetics compared to a no-coacervate control.** After lyophilization, reactions were rehydrated with 20 µL of lab-grade water and incubated at room temperature. Shading indicates the average value of n=3 independent biological replicates +/- standard deviation. Raw fluorescence values were standardized to µM FITC.

**Supplementary Data Files**

**Supplementary Data 1.** DNA (plasmids, oligonucleotides, and templates), RNA, and protein sequences used in this study.
**Supplementary Data 2.** Excel worksheet describing the assembly of ROSALIND reactions with spermine-PAA coacervates.
**Supplementary Data 3.** EC50, LOD, LOQ and VC values corresponding to each dose-response in this study.
**Supplementary Data 4.** Jupyter Notebook Python code used to calculate LOD, LOQ, and VC.

**Supplementary Information References**
