## Supplementary material for "Transcription-induced coacervation accelerates and sensitizes cell-free biosensing": Jupyter Notebook File: Feng_Transcription_Coacervation_SuppData4_SUBMIT.ipynb

```
# %% [code]
from scipy.optimize import brentq
import numpy as np

# functions are the Hill fits, and have the form f(x) = (A * x^B) / (C + x^B) + D
# define functions dictionary

FUNCTIONS = {
    # --- without coacervates series ---
    "f1": (0.983517,  7.13831,  0.11315,       0.0461971),
    "f2": (0.865166,  2.92932,  3.87881,       0.0139367),
    "f3": (0.971109,  2.86213,  1_096.33,      0.0332158),
    "f4": (0.833202,  9.30765,  1.78156e9,     0.0195293),
    "f5": (1.86404,   3.31183,  1.01991e6,     0.349355 ),
    "f6": (0.674783,  3.11148,  309_156,       0.102987 ),
    "f7": (0.892724,  2.49571,  71.3104,       0.125329 ),
    "f8": (0.782578,  4.36522,  8_765.23,      0.0791146),
    "f9": (0.53098,   3.98889,  13_011.3,      0.0173613),
    # --- with coacervates series ---
    "g1": (2.36249,   5.96813,  0.18367,       0.130053 ),
    "g2": (2.13391,   2.53806,  1.6468,        0.0142234),
    "g3": (2.19319,   1.85468,  50.2388,       0.109501 ),
    "g4": (1.60398,   5.84586,  449_468,       0.172495 ),
    "g5": (3.86024,   3.36116,  943_833,       0.680602 ),
    "g6": (1.76006,   2.95611,  47_431.8,      0.677734 ),
    "g7": (0.989978,  1.44268,  5.08402,       0.262859 ),
    "g8": (1.6575,    2.04428,  16.7543,       0.306225 ),
    "g9": (0.993506,  3.53018,  1_626.78,      0.100165 ),
}

# define large domain (0 to 10K) over which brentq will search for a solution 
X_MIN, X_MAX = 0, 10_000

# define functions
def f(x, A, B, C, D):
    xB = x ** B
    return (A * xB) / (C + xB) + D

# build inverse functions
def make_inverse(A, B, C, D, x_min=X_MIN, x_max=X_MAX): 
    def f_inv(y):
        return brentq(lambda x: f(x, A, B, C, D) - y, x_min, x_max)
    return f_inv

# create dictionary of inverse functions
inverses = {label: make_inverse(*params) for label, params in FUNCTIONS.items()}


# %% [code]
# The numbers plugged in below are the LOD thresholds, defined as mean(blank)+ 3 x std(blank) for each dose response.
# These LOD thresholds are also reported and detailed in Supplementary Data 3.
# The LOD is found by evaluating the inverse function at the LOD threshold for each dose response.

#print(inverses["f1"](0.041038588)) generates an error because D > 0.041
print(inverses["f2"](0.030257518))
print(inverses["f3"](0.053903238))
print(inverses["f4"](0.040871152))
print(inverses["f5"](0.357034409))
print(inverses["f6"](0.106737307))
print(inverses["f7"](0.197779321))
print(inverses["f8"](0.142317552))
print(inverses["f9"](0.107940982))

print("\n")

print(inverses["g1"](0.140191928))
print(inverses["g2"](0.066995754))
print(inverses["g3"](0.147305498))
#print(inverses["g4"](0.127922756)) generates an error because D > 0.13
#print(inverses["g5"](0.507923895)) generates an error because D > 0.51
print(inverses["g6"](0.763456932))
print(inverses["g7"](0.38630137))
print(inverses["g8"](0.441558977))
print(inverses["g9"](0.152278385))


# %% [code]
# The numbers plugged in below are the LOQ thresholds, defined as mean(blank)+ 10 x std(blank) for each dose response.
# These LOQ thresholds are also reported and detailed in Supplementary Data 3.
# The LOQ is found by evaluating the inverse function at the LOQ threshold for each dose response.

print(inverses["f1"](0.057952067))
print(inverses["f2"](0.061791929))
print(inverses["f3"](0.122853514))
print(inverses["f4"](0.084385321))
print(inverses["f5"](0.577126364))
print(inverses["f6"](0.162589604))
print(inverses["f7"](0.41563173))
print(inverses["f8"](0.324518678))
print(inverses["f9"](0.255389271))

print("\n")
print(inverses["g1"](0.242141537))
print(inverses["g2"](0.145896551))
print(inverses["g3"](0.297816906))
print(inverses["g4"](0.241021059))
# print(inverses["g5"](0.587853877)) -- errors because D > 0.59
print(inverses["g6"](1.134912249))
print(inverses["g7"](0.699543379))
print(inverses["g8"](0.78855529))
print(inverses["g9"](0.359852354))

# %% [code]
# The value "1" is plugged in below to determine the analyte concentration necessary to generate 1 μM FITC signal
# The VC is found by evaluating the inverse function at value 1

print(inverses["f1"](1))
#print(inverses["f2"](1)) generates an error because its asymptote (A+D) is 0.879
print(inverses["f3"](1))
#print(inverses["f4"](1)) generates an error because its asymptote (A+D) is 0.853
print(inverses["f5"](1))
#print(inverses["f6"](1)) generates an error because its asymptote (A+D) is 0.778
print(inverses["f7"](1))
#print(inverses["f8"](1)) generates an error because its asymptote (A+D) is 0.862
#print(inverses["f9"](1)) generates an error because its asymptote (A+D) is 0.548

print("\n")

print(inverses["g1"](1))
print(inverses["g2"](1))
print(inverses["g3"](1))
print(inverses["g4"](1)) 
print(inverses["g5"](1))
print(inverses["g6"](1))
print(inverses["g7"](1))
print(inverses["g8"](1))
print(inverses["g9"](1))
```
