## Supplementary material for "Transcription-induced coacervation accelerates and sensitizes cell-free biosensing": Source Data: Feng_Transcription_Coacervation_Image_SourceData_Fig.3_SUBMIT.docx

Feng, et al. (2026)

Source Data for Figure 3

The image below shows the uncropped and unprocessed Tris–acetate–EDTA–agarose gel image, showing two of the independent replicates quantified in **Figure 3B**.

Samples 2-4 are paired with samples 5-7. Samples 9-11 are paired with samples 12-14, and constitute a second, independent replicate.

From left to right:

1. Empty well
2. DNA, as-mixed before centrifugation
3. DNA with spermine-PAA coacervates, as-mixed before centrifugation
4. Spermine-PAA coacervates, as-mixed before centrifugation
5. DNA, supernatant collected after centrifugation
6. DNA with spermine-PAA coacervates, supernatant collected after centrifugation
7. Spermine-PAA coacervates, supernatant collected after centrifugation
8. Empty well
9. DNA, as-mixed before centrifugation
10. DNA with spermine-PAA coacervates, as-mixed before centrifugation
11. Spermine-PAA coacervates, as mixed before centrifugation
12. DNA, supernatant collected after centrifugation
13. DNA with spermine-PAA coacervates, supernatant collected after centrifugation
14. Spermine-PAA coacervates, supernatant collected after centrifugation
15. Empty well

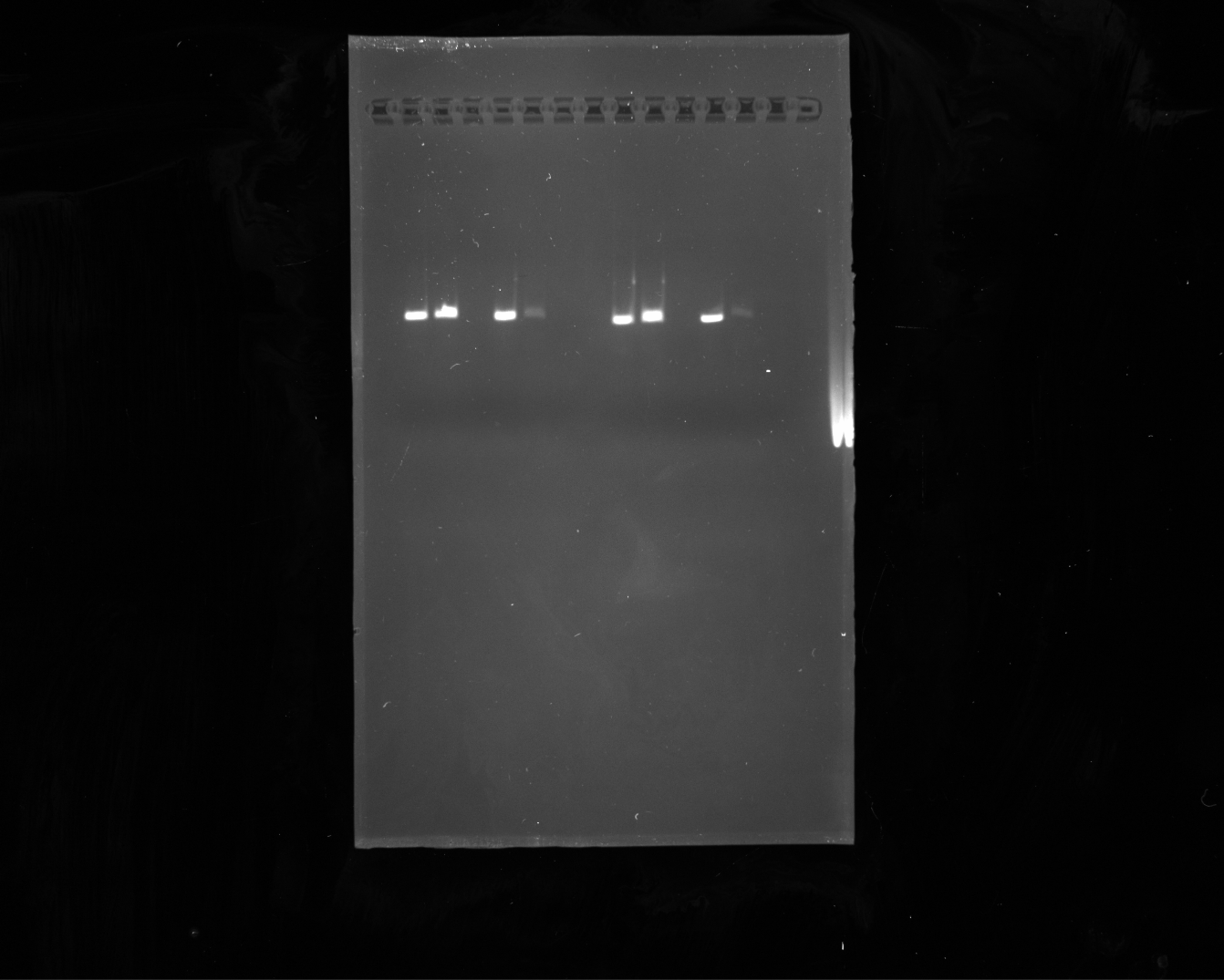

10 11 12 13 14 15

1	2	3	4	5	6

7	8	9

The image below shows the uncropped and unprocessed Tris–acetate–EDTA–agarose gel image, showing a third independent replicate quantified in **Figure 3B.**

From left to right:

1. Empty well
2. DNA, as-mixed before centrifugation
3. DNA with spermine-PAA coacervates, as-mixed before centrifugation
4. Spermine-PAA coacervates, as-mixed before centrifugation
5. DNA, supernatant collected after centrifugation
6. DNA with spermine-PAA coacervates, supernatant collected after centrifugation
7. Spermine-PAA coacervates, supernatant collected after centrifugation
8. Empty well
9. Empty well
10. Empty well
11. Empty well
12. Empty well
13. Empty well
14. Empty well
15. Empty well

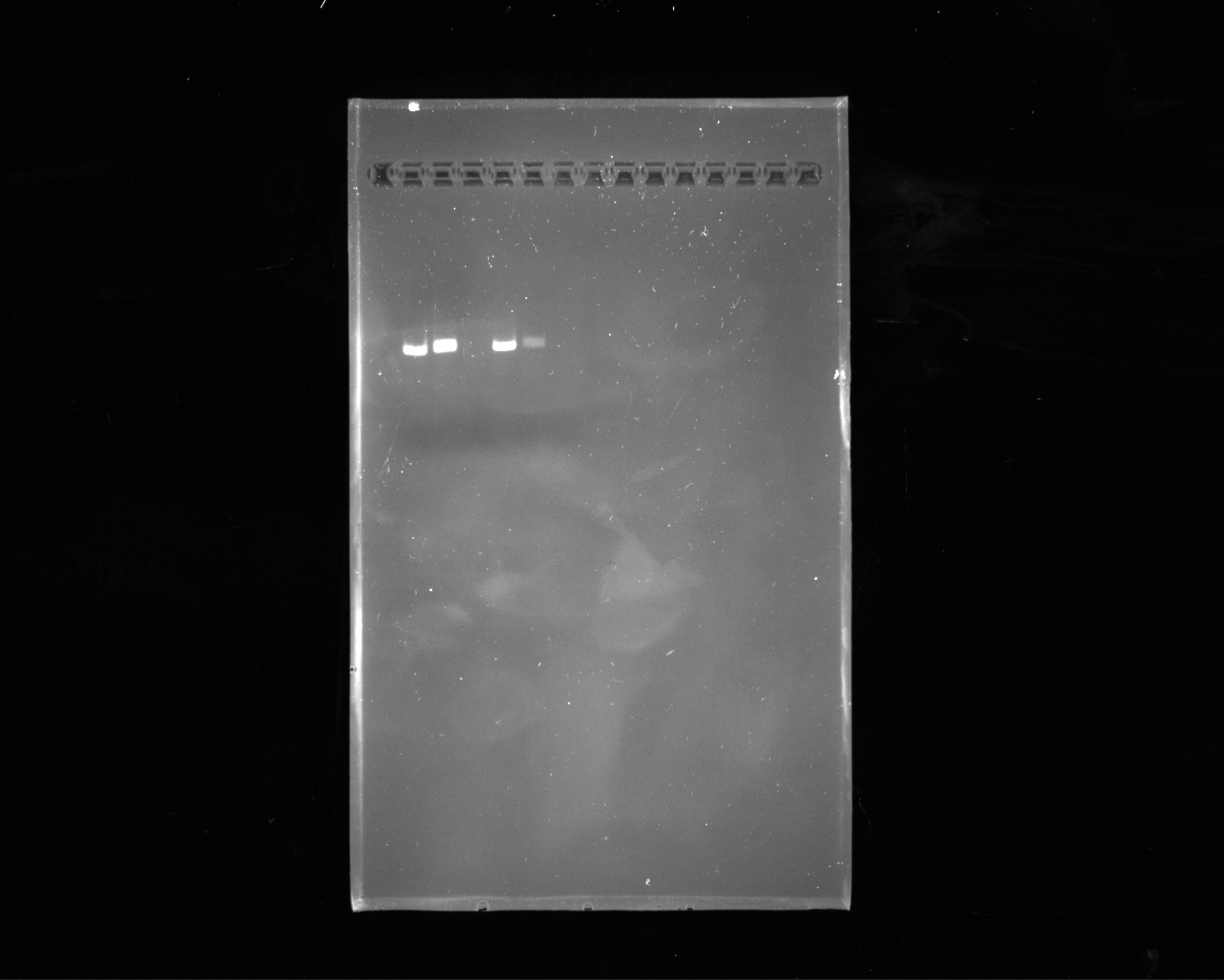

1	2	3	4	5

6	7	8	9

10 11 12 13 14 15
