## Supplementary material for "Transcription-induced coacervation accelerates and sensitizes cell-free biosensing": Source Data: Feng_Transcription_Coacervation_Image_SourceData_SI_Fig.4_SUBMIT.docx

Feng, et al. (2026)

Source Data for Supplementary Figure 4

The image below shows the uncropped and unprocessed urea-PAGE gel image quantified in **Supplementary Figure 4**.

From left to right:

1. Empty well
2. Empty well
3. Century-Plus RNA Ladder (1000, 750, 500, 400, 300, 200 and 100 nts).
4. IVT assembled without coacervates (replicate 1)
5. IVT assembled with coacervates (replicate 1)
6. IVT assembled without coacervates (replicate 2)
7. IVT assembled with coacervates (replicate 2)
8. IVT assembled without coacervates (replicate 3)
9. IVT assembled with coacervates (replicate 3)
10. Empty well

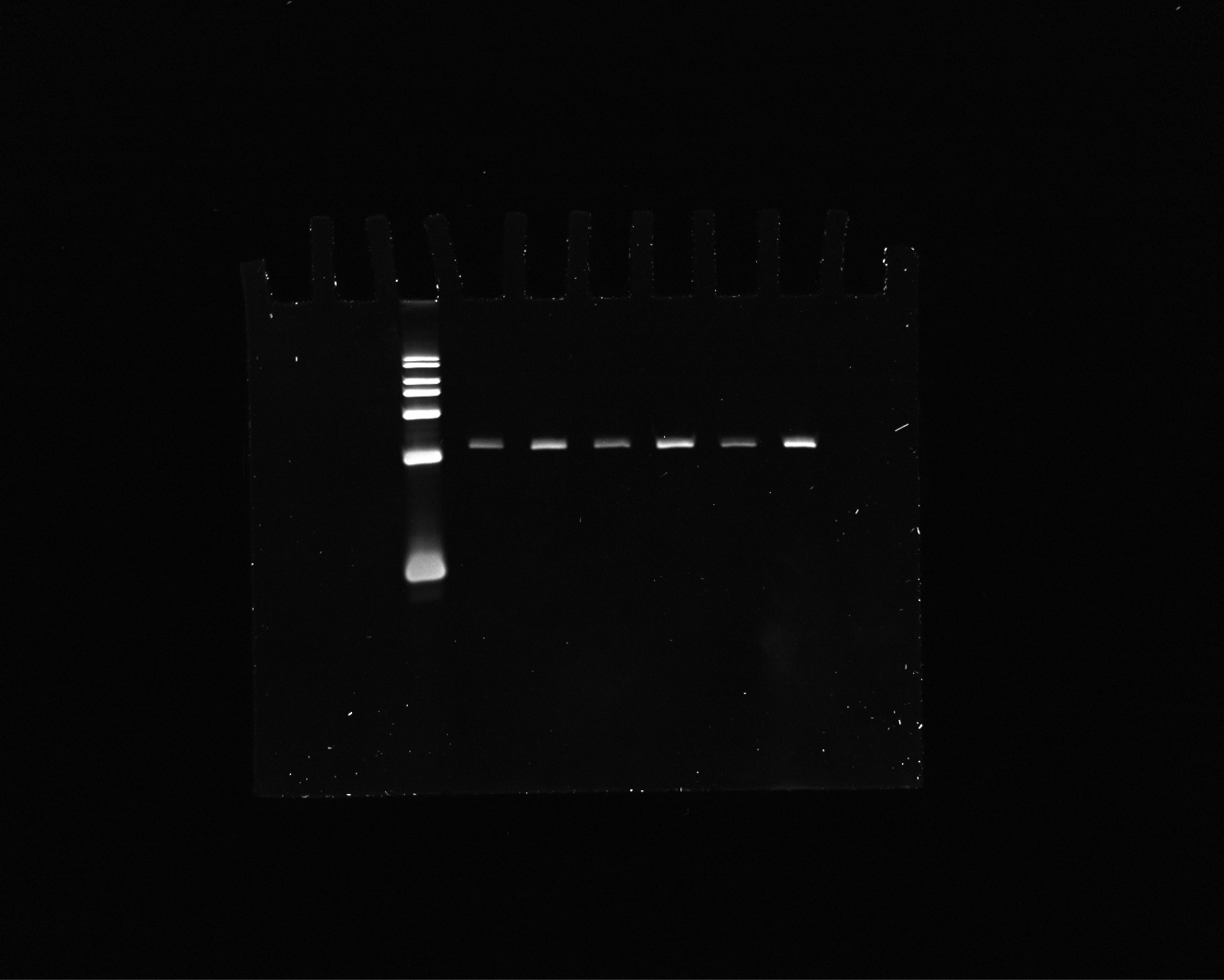

9

8

7

5

10

3

6

4

1

2

- 4
