## Supplementary material for "Transcription-induced coacervation accelerates and sensitizes cell-free biosensing": Source Data: Feng_Transcription_Coacervation_Image_SourceData_SI_Fig.5_SUBMIT.docx

Feng, et al. (2026)

Source Data for Supplementary Figure 5

The image below shows the uncropped and unprocessed image of tubes shown in **Supplementary Figure 5B**.

Tubes from left to right:

1. DNA, as-mixed before centrifugation
2. DNA with spermine-PAA coacervates, as-mixed before centrifugation

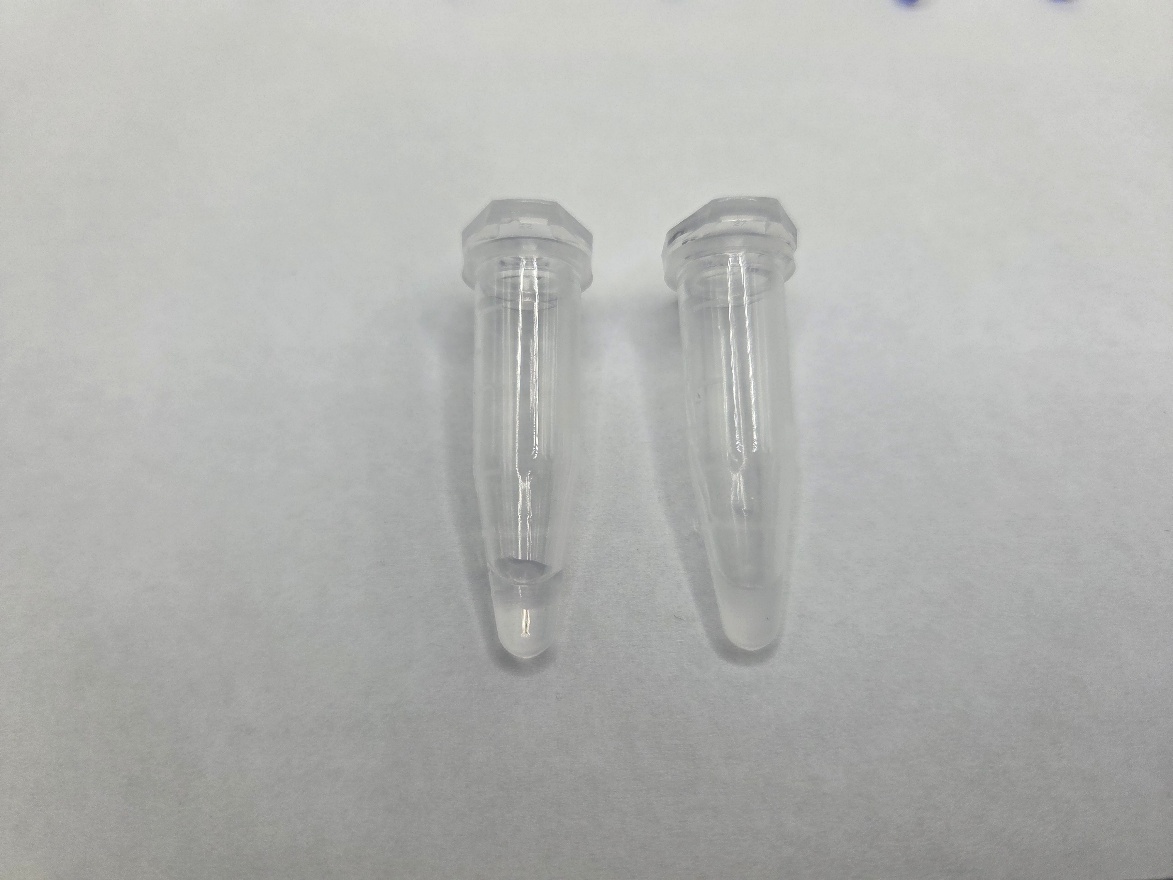

The image below shows the uncropped and unprocessed image of tubes shown in **Supplementary Figure 5C**.

Tubes from left to right:

1. DNA, as-mixed after centrifugation
2. DNA with spermine-PAA coacervates, as-mixed after centrifugation

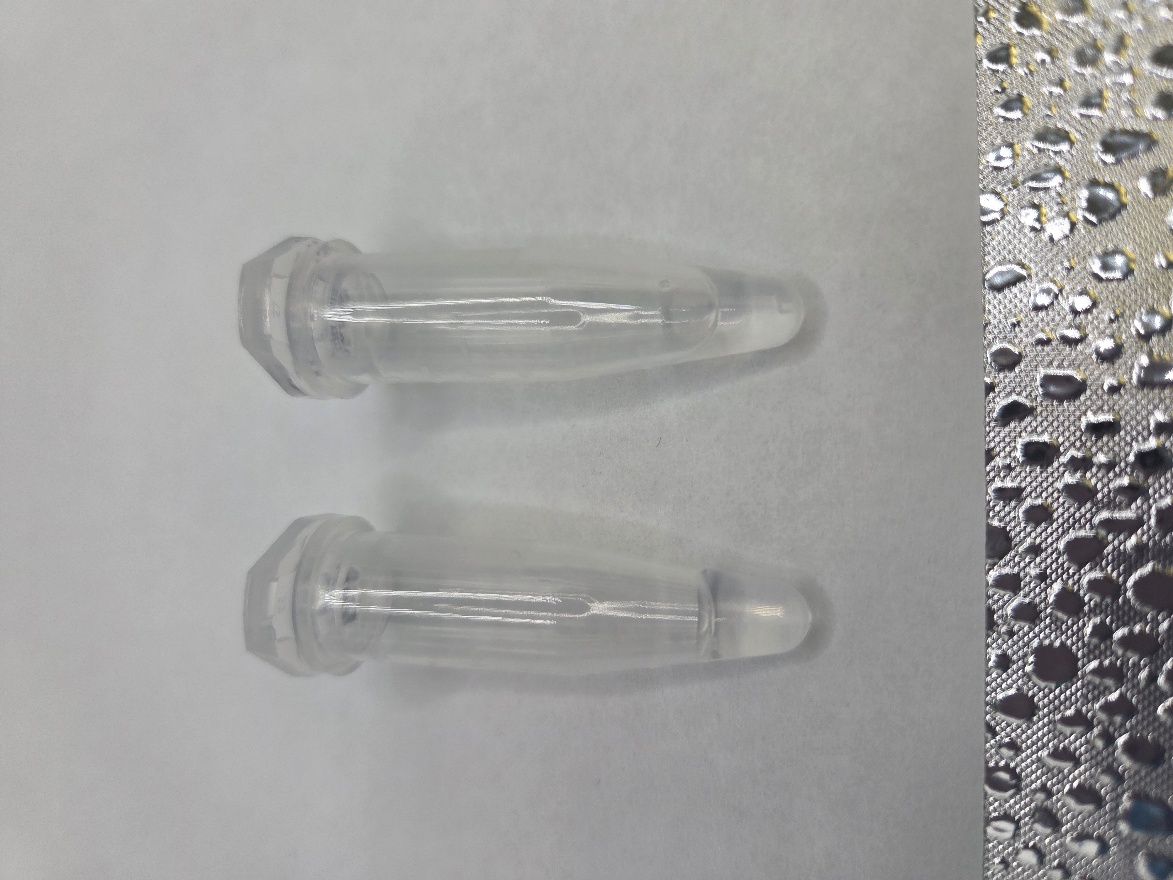
