## Supplementary material for "Transcription-induced coacervation accelerates and sensitizes cell-free biosensing": Source Data: Feng_Transcription_Coacervation_Image_SourceData_SI_Fig.7_SUBMIT.docx

Feng, et al. (2026)

Source Data for Supplementary Figure 7

The image below shows the uncropped and unprocessed image of tubes shown in **Supplementary Figure 7A-B**.

Tubes from left to right:

1. DNA with spermine-PAA coacervates, after centrifugation
2. 0.5 µL of 0.024% bromophenol blue in formamide
3. 1 µL of 0.024% bromophenol blue in formamide
4. 2 µL of 0.024% bromophenol blue in formamide
5. 3 µL of 0.024% bromophenol blue in formamide

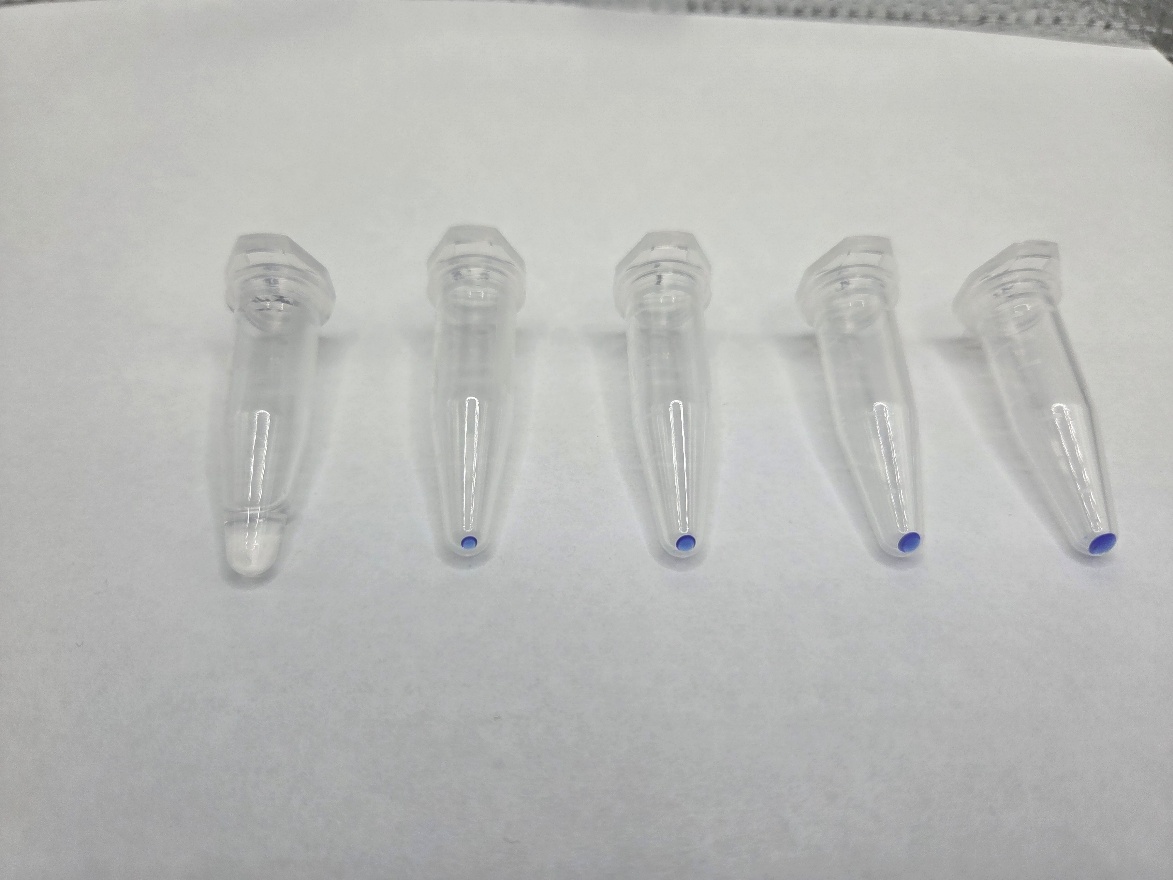
