## Supplementary material for "Transcription-induced coacervation accelerates and sensitizes cell-free biosensing": Source Data: Feng_Transcription_Coacervation_Image_SourceData_SI_Fig.12_SUBMIT.docx

Feng, et al. (2026)

Source Data for Supplementary Figure 12

The image below shows the uncropped and unprocessed image of tubes shown in **Supplementary Figure 12**

The photos are arranged as they appear in Supplementary Figure 12. In clockwise direction, starting from the top left:

1. Biosensor reactions with coacervates illuminated under room light
2. Biosensor reactions with coacervates after 1 hour of room temperature incubation, illuminated via blue light LEDs
3. Biosensor reactions with coacervates after 2 hours of room temperature incubation, illuminated via blue light LEDs
4. Biosensor reactions without coacervates after 2 hours of room temperature incubation, illuminated via blue light LEDs
5. Biosensor reactions without coacervates after 1 hour of room temperature incubation, illuminated via blue light LEDs
6. Biosensor reactions without coacervates illuminated under room light

In each photo, the tube on the left contains 0 µM anhydrotetracycline, and the tube on the right contains 1.9 µM anhydrotetracycline.

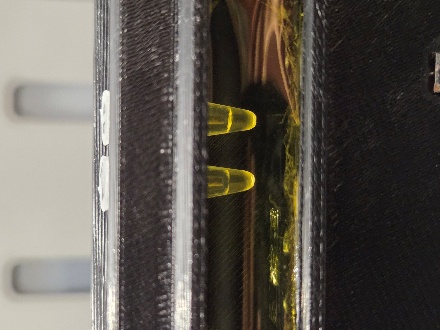

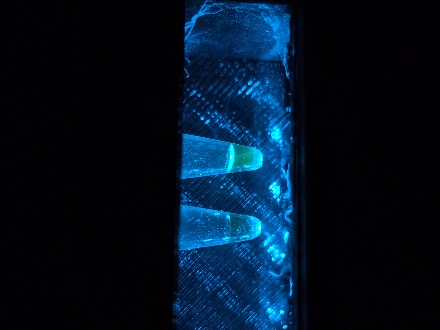

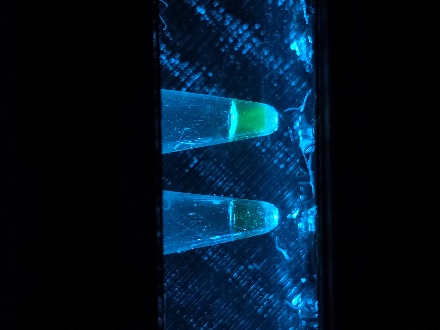

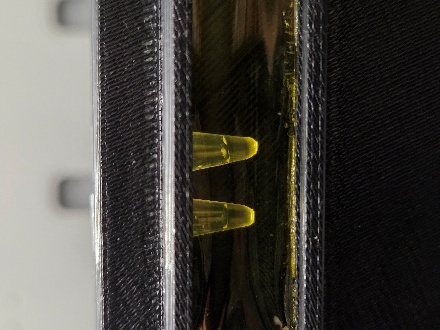

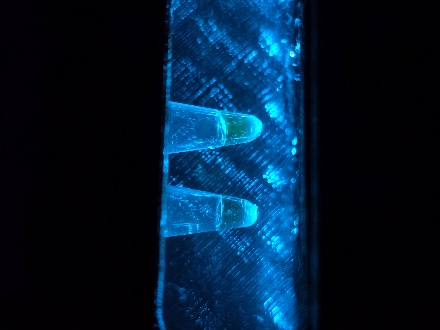

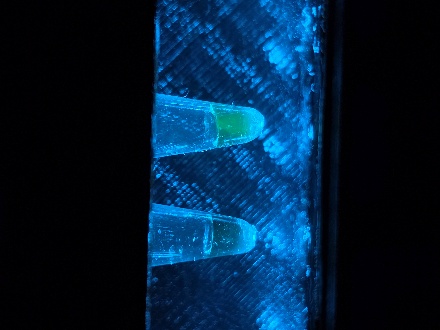
